# Microglia control small vessel calcification via TREM2

**DOI:** 10.1101/829341

**Authors:** Yvette Zarb, Sina Nassiri, Sebastian Guido Utz, Johanna Schaffenrath, Elisabeth J. Rushing, K. Peter R. Nilsson, Mauro Delorenzi, Marco Colonna, Melanie Greter, Annika Keller

## Abstract

Microglia participate in CNS development and homeostasis and are often implicated in modulating disease processes in the CNS. However, less is known about the role of microglia in the biology of the neurovascular unit (NVU). In particular, data are scant on whether microglia are involved in CNS vascular pathology. In this study, we use a mouse model of primary familial brain calcification (PFBC) – *Pdgfb^ret/ret^* to investigate the role of microglia in calcification of the NVU. We report that microglia enclosing vessel-calcifications, coined calcification-associated microglia (CAM), display a distinct activation signature. Pharmacological ablation of microglia with the CSF1R inhibitor - PLX5622 leads to aggravated vessel calcification. Additionally, depletion of microglia in wild-type and *Pdgfb^ret/ret^* mice causes the development of bone protein (osteocalcin, osteopontin) containing axonal spheroids in the white matter. Mechanistically, we show that microglia require functional TREM2 for controlling vessel-associated calcification. In conclusion, our results demonstrate that microglial activity in the setting of pathological vascular calcification is beneficial. In addition, we identify a new, previously unrecognized function of microglia in halting the expansion of ectopic calcification.

## Introduction

Microglia participate in CNS development and homeostasis by regulating neural cell numbers, migration of interneurons as well as promoting connectivity, synapse formation and pruning ^1^. Microglia are considered the first line of immune defence in the brain by monitoring brain parenchyma under homeostatic conditions and resolving cerebral insults ^2, 3^. However, less is known about the function of microglia at the neurovascular unit (NVU), which is composed of vascular cells (endothelial and mural cells, as well as perivascular fibroblasts and macrophages), associated neurons, and glia ^4^. Microglia have been implicated in shaping brain vasculature by inducing anastomosis of vascular sprouts ^5^. In adults, microglial processes directly contact the endothelial basement membrane at the NVU ^6^; however, the functional importance of these contacts is poorly understood. Microglia protect against intrinsic (e.g. aggregated, mutated proteins, tumour cells) and extrinsic (e.g. pathogens) injury ^7^. During the course of brain diseases, microglia lose the homeostatic signature and acquire a neurodegenerative signature that is partially shared by diseases such as Alzheimer’s disease (AD), multiple sclerosis and amyotrophic lateral sclerosis ^8–10^. Disruption of the NVU is common to many neurodegenerative diseases ^4^, but studies on microglial function have mainly focused on brain parenchyma and not the NVU. In normal mouse brain, laser-induced vascular injury results in microglial activation with the rapid development of processes that shield a lesioned blood vessel section and phagocytose debris ^2^. These findings support the important role of microglia in vascular repair.

Optimal functioning of the NVU, which mediates hyperaemia, is crucial for cerebral perfusion ^4^. Blood vessels play an integral role in brain development and provide a niche for brain stem cells. In addition, cerebral vasculature senses the environment and communicates changes to neural tissue, participates in glymphatic clearance, and controls immune quiescence in the CNS ^11–15^. Accordingly, dysfunction of the NVU accompanies or may even represent a primary cause of many neurodegenerative diseases ^4, 16^. In the case of the primary familial brain calcification (PFBC), bilateral basal ganglia calcification of blood vessels is a key diagnostic criterion. The pathogenic mechanism points to a compromised NVU ^17–20^. PFBC is a clinically and genetically heterogenous disease caused by mutations in at least in five genes – *MYORG, PDGFB, PDGFRB, SLC20A2* and *XPR1* ^21–25^. Of note, recent studies have estimated the minimal prevalence of PFBC ranges from 4-6 p. 10,000, depending of the causative gene mutation, thus suggesting that PFBC is not a rare disorder and is likely underdiagnosed ^26, 27^. In addition, basal ganglia calcification is a common radiological finding, estimated in up to 20% of patients undergoing CT imaging ^28, 29^. Although the effect of cerebral calcification on the NVU and brain parenchyma is unknown, peripheral vascular calcification can lead to cardiovascular morbidity and mortality ^30^.

Leukodystrophies and interferonopathies linked to microglial dysfunction are associated with brain calcifications. In addition to cerebral atrophy, patients with Nasu-Hakola disease, caused by mutations in triggering receptor expressed on myeloid cell 2 (*TREM2*) ^31^ and TYRO protein tyrosine kinase binding protein (*TYROBP*) ^32^ exhibit vascular calcification in the basal ganglia ^33–35^. Furthermore, mice lacking *Usp18* in microglia, a protein negatively regulating interferon signalling, develop basal ganglia calcification ^36^. Additionally, *USP18* deficiency was reported to cause type 1 interferonopathy, a group of monogenic autoimmune disorders presenting frequently with cerebral calcification ^37^. Interestingly, patients with mutations in other genes implicated in microglial development and function (e.g. *IRF8*, *CSF1R*) develop intracerebral calcifications ^38–40^. Thus, cell-autonomous defects in microglia lead to the formation of brain calcifications. The precise role of microglia in the pathology of brain vascular calcifications remains to be determined.

In this study, we investigated the role of microglia in vascular calcification using a mouse model of PFBC. Previously, we described that mouse platelet-derived growth factor B (*Pdgfb*) hypomorphs (*Pdgfb^ret/ret^*) develop brain vessel-associated calcifications similar to human PFBC ^22^. Vascular calcification in human PFBC and mouse models of PFBC were conspicuously encircled by activated microglia ^18, 19, 22, 41, 42^ and demonstrated an osteogenic environment with the surrounding cells expressing osteoclast markers ^18^. This observation raised the question of the cellular origin and function of osteoclast-like cells. Another aspect to be clarified is whether microglia participate in the development of NVU calcification in PFBC.

We characterize transcriptome changes in calcified tissue in *Pdgfb^ret/ret^* mice and demonstrate that microglia encircling vascular calcifications, coined “calcification-associated microglia” (CAM), express a distinct activation signature. Pharmacological ablation of microglia in *Pdgfb^ret/ret^* mice leads to aggravated calcification of the NVU. Additionally, we show in wild-type mice and a mouse model of PFBC that ablation of microglia leads to the development of axonal spheroids containing bone proteins (osteocalcin, osteopontin) in the white matter. Mechanistically, we show that microglia require functional TREM2 for controlling vessel-associated calcifications, as genetic deletion of *Trem2* in *Pdgfb^ret/ret^* mice exacerbates calcification. In conclusion, our study shows for the first time that microglia play an important role in modifying vascular calcification in the brain, and identifies microglia as a potential therapeutic target in PFBC.

## Materials and Methods

### Mice

Mice used in this study were 1 to 5 months old. The following mouse strains were used: C57BL6/J (Charles River), B6.Pdgfb^tm3Cbet^ (*Pdgfb^ret/ret^*) ^22, 43^, *Cx3cr1-*CreER^T2 44^, *Sall1-* CreER^T2 45^, B6.Cg-Gt(ROSA)26Sor^tm14^(CAG–tdTomato)^Hze^/J (Ai14) (Jackson Stock # 007914), APP/PS1 ^46^ and *Trem2^-/-^* ^47^. This study was carried out in accordance with study protocols approved by the Cantonal Veterinary Office Zurich (permit numbers: ZH196/2014, ZH151/2017).

### In vivo treatments

#### Tamoxifen treatment for the genetic labelling of Cx3Cr1 and Sall1 -expressing cells

For tamoxifen-inducible labelling of *Cx3cr1* and *Sall1* expressing cells in respective genotypes (*Cx3cr1-*CreER^T2^; Ai14^Tg/wt^; *Pdgfb^ret/ret^* or *Pdgfb^ret/wt^* and *Sall1-*CreER^T2^; Ai14^Tg/wt^; *Pdgfb^ret/ret^* or *Pdgfb^ret/wt^*), 4 week-old animals were treated for five consecutive days via oral gavage with 2 mg of tamoxifen (Sigma-Aldrich, cat # T5648) dissolved in corn oil. Mice were sacrificed at 4 months of age.

#### Pharmacological ablation of microglia

Oral CSF1R inhibitor, PLX5622 (Plexxikon Inc.) ^48, 49^, was formulated in AIN-76A standard chow by Research Diets (New Brunswick, NJ) at 1200 ppm. Control mice received AIN-76A chow without PLX5622. PLX5622 chow and control chow were provided by Plexxikon, Inc. under a Materials Transfer Agreement. One-month-old mice were fed with chow containing PLX5622 or control chow for 2 months and sacrificed at the age of 3 months.

#### Edu treatment

Three-month-old mice were injected with Edu (50 mg/kg, Sigma-Aldrich, cat # 900584) intra-peritoneally for three consecutive days and sacrificed on the following day.

### Antibodies

Primary antibodies used for immunofluorescence staining are listed in Supplementary Table 1. All secondary antibodies (suitable for multiple labelling) labelled with various fluorophores (Alexa 488, Cy3, DyLight 649) made in donkey (anti-rabbit, anti-rat and anti-goat) or in goat (anti-chicken Cy3) were purchased from Jackson Immunoresearch. Antibodies used for flow cytometry analysis are listed in Supplementary Table 2.

### Histochemistry and immunohistochemistry

Immunohistochemistry was performed according to methods described previously ^18^. For Edu detection, the Click iT™ Edu Alexa Fluor™ 555 imaging kit (Thermo Fischer) was used and slices were treated according to the manufacturer’s instructions. Immunohistochemical stainings were imaged with a confocal microscope (Leica SP5, 20x numerical aperture (NA): 0.7, 40x NA: 1.25, 63x NA: 1.4) or stereomicroscope (Zeiss Axio Zoom.V16, 1x NA: 0.25). For stains that exhibited salt-and-pepper noise, a median filter of 5 x 5 x 5 was applied to eliminate noise. Images were analysed using the image-processing software Imaris 9.2.0. (Bitplane) and Adobe Illustrator CS6.

For histochemistry, mouse brains were collected and embedded in paraffin. Two µm thick tissue sections were stained with periodic acid-Schiff (PAS) or the Bielschowsky silver stain using standard protocols. For alizarin red staining, sections were deparaffinised and rehydrated, incubated for 1 hour in 1% alizarin red solution (pH 9.0) followed by 1 hour in 1% alizarin red solution (pH 6.4) at room temperature. Stained paraffin sections were scanned with NanoZoomer HT (Hamamatsu Photonics), equipped with a 20× objective (UPlanSapo, NA: 0.75, Olympus). Images were analysed using Digital Image Hub software (SlidePath) and Adobe Illustrator CS6.

### Detection of aggregated proteins

Paraffin embedded 6µm thick brain sections were used from *Pdgfb^ret/ret^* and controls of 1 year of age or older. As a positive control, paraffin embedded sections from APP/PS1 mice ^46^ were used. Sections were deparaffinized and hydrated, followed by a 30-minute incubation with luminescent-conjugated oligothiophene (LCO) h-HTAA ^50^ at room temperature. For Thioflavin T (Sigma Aldrich, T3516) staining, deparaffinized sections were processed according to the manufacturer’s instructions. Congo red staining was performed according to standard protocol. Stained sections were imaged with a confocal microscope (Leica SP5, 20x, NA: 0.7) or Axioplant (Zeiss, 20x, NA: 0.5). Images were analyzed with image-processing software Imaris (Bitplane).

### Quantification of immunofluorescent stainings

Images for quantification of brain calcifications, cathepsin K intensity and astrocyte reactivity were acquired using a 20x objective (NA: 0.7, Leica SP5) 42 z-stacks with a 1.48µm step, 512×512 pixel resolution. For quantification of total brain calcification, z-stacks of APP staining and osteocalcin were summed in Fiji (ImageJ) ^51^. Imaris software was used to quantify calcification using the function “surfaces”. Quantification of calcifications was performed in mid-midbrain, which shows the smallest inter-individual variation in calcification load ^18^. Cathepsin K intensity density was calculated using Fiji (ImageJ) software. Astrocyte reactivity was quantified using GFAP, podoplanin, LCN2, C3 staining in Fiji (ImageJ). The LCN2 and C3 signal intensity was first masked to GFAP staining to eliminate non-astrocytic expression/deposition. Signal intensity was then normalized to the GFAP signal. Two technical replicates were quantified for each animal.

Microglia were quantified on images acquired using a 20x objective (NA: 0.7, Leica SP5), 41 z-stacks with a 0.64 µm step, 1024×1024 pixel resolution. Quantification of microglia was performed using the morpholibJ package ^52^ in Fiji, with minor modifications as described previously ^53^.

### Flow cytometry analysis

Mice were deeply anaesthetized using a mixture of ketamine and xylazine and perfused transcardially using ice-cold PBS. Subsequently, mouse brains were dissected into non-calcification-prone brain regions, i.e., cortex, hippocampus and cerebellum, and calcification-prone regions, i.e., thalamus, midbrain and pons. Brain cell suspensions were prepared by cutting the tissue into small pieces, followed by collagenase type IV treatment. Dissociated tissue was passed through an 18G syringe to obtain a homogeneous cell suspension and further enriched with a Percoll gradient. Samples were then passed through a 70 µm filter, followed by red blood cell lysis and antibody staining. Flow cytometric analysis was carried using FACSymphony (BD Biosciences) and analysed with FlowJo and R software.

### High-dimensional analysis

Raw data was pre-analysed with FlowJo, subsequently transformed in Matlab using cyt3 and percentile normalized in R. Dimensionality reduction was achieved by uniform manifold approximation and projection (UMAP). FlowSOM was used for automated and expert-guided cell clustering ^54^. Mean marker expression was projected onto UMAP in order to generate a heatmap of median expression values ^55^.

### Isolation of vessel calcifications and RNA sequencing

Mice were deeply anaesthetized and transcardially perfused with ice-cold PBS. Brains were removed, placed in RNA*later*™ Stabilization Solution (Thermo Fischer Scientific, Cat # AM7020) and 1 mm coronal sections were cut using a brain matrix (RBMA-200C, World Precision Instruments). Calcifications were detected based on their auto-fluorescence ^22^ using a fluorescent stereomicroscope (Zeiss Axio Zoom.V16) and were surgically removed together with surrounding tissue. Cortical sections were also removed as examples of non-calcification prone regions. RNA was isolated with a micro RNA kit (Qiagen) according to the manufacturer’s instructions. The concentration of RNA and sample purity were assessed using a 2100 Bioanalyzer (Agilent) and RNA 6000 Pico Kit (Agilent). RNA samples were polyA-enriched, and libraries were prepared using the Illumina TruSeq® Stranded RNA kit. RNA was sequenced on an Illumina platform Hi-Seq 4000 at the Functional Genomic Centre Zurich (UZH, ETH). The Illumina single read approach (1×125 bp) was used to generate raw sequencing reads with a depth of 20–30 million reads per sample.

### Bioinformatics analysis

Quantification of RNA-seq data was performed using kallisto ^56^. In brief, target transcript sequences were obtained from ENSEMBLE (GRCm38.p6), and the abundance of transcripts was quantified using kallisto 0.44.0 with sequence-based bias correction. All other parameters were set to default when running kallisto. Kallisto’s transcript-level estimates were further summarized at the gene-level using tximport 1.8.0 from Bioconductor ^57^.

For downstream analysis, genes of low abundance were filtered out and unwanted variation was estimated using the RUVr functionality from the RUVseq 1.16.0 package within Bioconductor ^58^. The number of factors of unwanted variation estimated from the data was set to 3, and the genes_by_samples matrix of residuals was obtained from a first-pass quasi-likelihood negative binomial generalized log-linear regression of the counts on biological covariates using the edgeR package from Bioconductor ^59^.

Differential expression analysis was performed using DESeq2 1.22.0 from Bioconductor ^60^, with estimated factors of unwanted variation included as additional covariates in the design formula. Significant genes were identified using FDR<0.05 and foldchange>2.

Co-expression network analysis was performed using WGCNA R package. Briefly, normalized count data obtained from RUVseq were first adjusted for mean-variance trend using the regularized log transformation of DESeq2 1.22.0 from Bioconductor ^60^. To exclude uninformative genes from co-expression analysis in an unbiased manner, 867 highly variable genes were selected based on the overall distribution of coefficients of variation. Subsequently, a signed weighted network of highly variable genes was constructed using biweight midcorrelation with beta=23 as soft thresholding power. Hierarchical clustering of the topological overlap matrix dissimilarity further revealed 6 modules of positively correlated genes. In order to identify hub genes associated with brain calcification, we computed module membership and gene significance for each gene belonging to the module associated with brain calcification.

Gene Set Enrichment Analysis (GSEA) was performed using fgsea 1.8.0 package from Bioconductor ^61^ with signal to noise ratio as defined by ^62^ as gene-level statistic. Prior to GSEA, mouse genes were converted to human orthologs using biomaRt 2.38.0 from Bioconductor ^63^. If a human ortholog was associated with more than one mouse gene, the mouse gene with maximum mean expression was selected using the collapseRows functionality within the WGCNA R package ^64^. Signaling pathways analyzed by GSEA were obtained from the Hallmark gene sets of the MSigDB ^65^. Gene signature of disease-associated microglia (DAM and proliferative-region-associated microglia (PAM) were obtained from literature ^8, 66^. Heatmaps were generated using the pheatmap R Package ^67^, with clustering distance and method set to Euclidean and ward.D2, respectively.

RNA sequencing data, both raw data and gene-by-sample matrix of estimated counts were deposited in Gene Expression Omnibus (GEO) under accession number GSE135449.

### Statistical analysis

Quantified values are represented as mean ±SEM. The following statistical tests were performed with Prism8 software (GraphPad). Normality was assessed using a Shapiro-Wilk test. Following tests were used to calculate statistical significance: student’s t-test (unpaired, two tailed), one-way ANOVA with Dunnett’s multiple comparison or Mann-Whitney two-tailed test. P-values < 0.05 were considered significant.

## Results

### Brain vessel-associated calcifications trigger an inflammatory environment

Vascular calcification in PFBC elicits a conspicuous glial reaction ^18, 19, 22, 41^. In order to gain insights into inflammatory changes accompanying vessel calcifications, we performed transcriptome analysis using RNA sequencing (RNA-seq) on tissue enriched with vessel-associated calcifications isolated from brains of *Pdgfb^ret/ret^* mice - a mouse model of PFBC. Brain calcifications are autofluorescent ^22^, which we exploited to isolate calcifications manually from brains of *Pdgfb^ret/ret^* animals under a fluorescent stereomicroscope. Non-calcified tissue from the same anatomical region (thalamus/midbrain) was also collected from control animals (*Pdgfb^ret/wt^*). This brain region is referred to as a “calcification-prone region” (Fig. 1A). We also isolated tissue from the cortex (referred to as a “non-calcification-prone region”) of both *Pdgfb^ret/ret^* and control animals (Fig. 1A). Principal component analysis (PCA) of transcriptomic data showed that the first PC accounts for variability due to anatomical differences (thalamus/midbrain vs. cortex), while the second PC accounts for differences between genotypes (Suppl Fig. 1A). When comparing calcification- and non-calcification-prone brain regions of *Pdgfb^ret/ret^* and control animals, we detected 92 and 94 deregulated genes, respectively (false discovery rate (FDR) <0.05 and fold change >2, Fig. 1B, Suppl Fig. 1B, Suppl Table 3 and 4). Contrasting differentially expressed genes (DEGs) in non-calcification prone and calcification-prone region between *Pdgfb^ret/ret^* and control animals showed that 74 genes were deregulated only in a calcification-prone region (thalamus/midbrain), and 60 genes were deregulated only in a non-calcification-prone region (cortex) (Fig. 1C, D, Suppl. table 5). The remaining 26 deregulated genes detected in both regions (Fig. 1C, D, Suppl. table 5) concur with previously reported vascular transcriptional alterations due to reduced pericyte numbers in *Pdgfb^ret/ret^* mice ^68^. Enrichment analysis of hallmark pathways showed that several inflammatory pathways, such as interferon gamma and alpha pathways, the reactive oxygen species pathway, and the inflammatory response pathway are enriched in *Pdgfb^ret/ret^* mice compared to controls (Fig. 1E). Several pathways were enriched only in calcification-prone regions, such as IL2-STAT5 signalling, unfolded protein response and TNFA signalling via NF-κb (Suppl Fig. 1C). Various significantly deregulated hallmark pathways (Fig. 1E, Suppl Fig. 1C) have been associated with activated microglia (e.g. interferon-related, complement-related) or microglia quiescence (e.g. TGF-beta-related) ^69, 70^. Notably, *Cst7*, which encodes a cysteine protease inhibitor and expressed in activated microglia ^8, 71, 72^, was the top upregulated gene (3.7 fold) in brain regions with calcifications (Fig. 1B, Suppl Table 3). The expression of *Cst7* was not detected in either genotype in a non-calcification-prone brain region (cortex) or in the calcification-prone region of control animals (Suppl Fig. 1D, Suppl Table 3, 4). To further explore the gene signature associated with vascular calcification, we performed co-expression network analysis on highly variable genes in the RNA-seq dataset (Suppl Fig. 1E) and identified six modules of positively correlated genes (Suppl Fig. 1F, Suppl Table 6). We found that only one module (turquoise) was associated with the calcified brain region in the *Pdgfb^ret/ret^* mice (Fig. 1F). Interestingly, among the hub genes within the turquoise module we identified *Cst7* and *Itgax* (Fig. 1G), genes induced in microglia in neurodegenerative diseases - termed DAM (disease-associated microglia) and in aging ^8, 71^. Furthermore, in addition to *Cst7* and *Itgax*, several other genes associated to reactive microglia or the DAM signature (e.g. *Cd68, Clec7a, Lpl*) were significantly upregulated in calcified brain regions of *Pdgfb^ret/ret^* mice (Fig. 1B, 2A). However, when comparing upregulated genes in calcification-prone regions in *Pdgfb^ret/ret^* mice using stringent criteria (FDR <0.05 and fold change >2) with the DAM signature, we found an overlap of only six genes (*Lpl, Cd68, Tyrobp, Itgax, Cst7, Clec7a*) (Suppl Fig. 1G). The DAM signature was recently reported to overlap with the “proliferative-region associated microglia” (PAM) signature that defines a subset of microglia found in white matter during development ^66^. By comparing the PAM signature with our dataset using the same criteria as for the DAM genes, an overlap of seven genes (*Hmox1, Lpl, Cd68, Slc16a3, Lag3, Tyrobp, Clec7a*) was found (Suppl Fig. 1H). We validated the expression of DAM/PAM-signature genes TIMP2 (Fig. 2B), CD68 (Fig. 2C) and CLEC7A (Fig. 2D) in microglia surrounding vessel-associated calcifications with immunohistochemistry. Thus, CAM exhibited an activation profile overlapping with microglia during aging, neurodegenerative proteinopathies and development.

**Figure 1.**
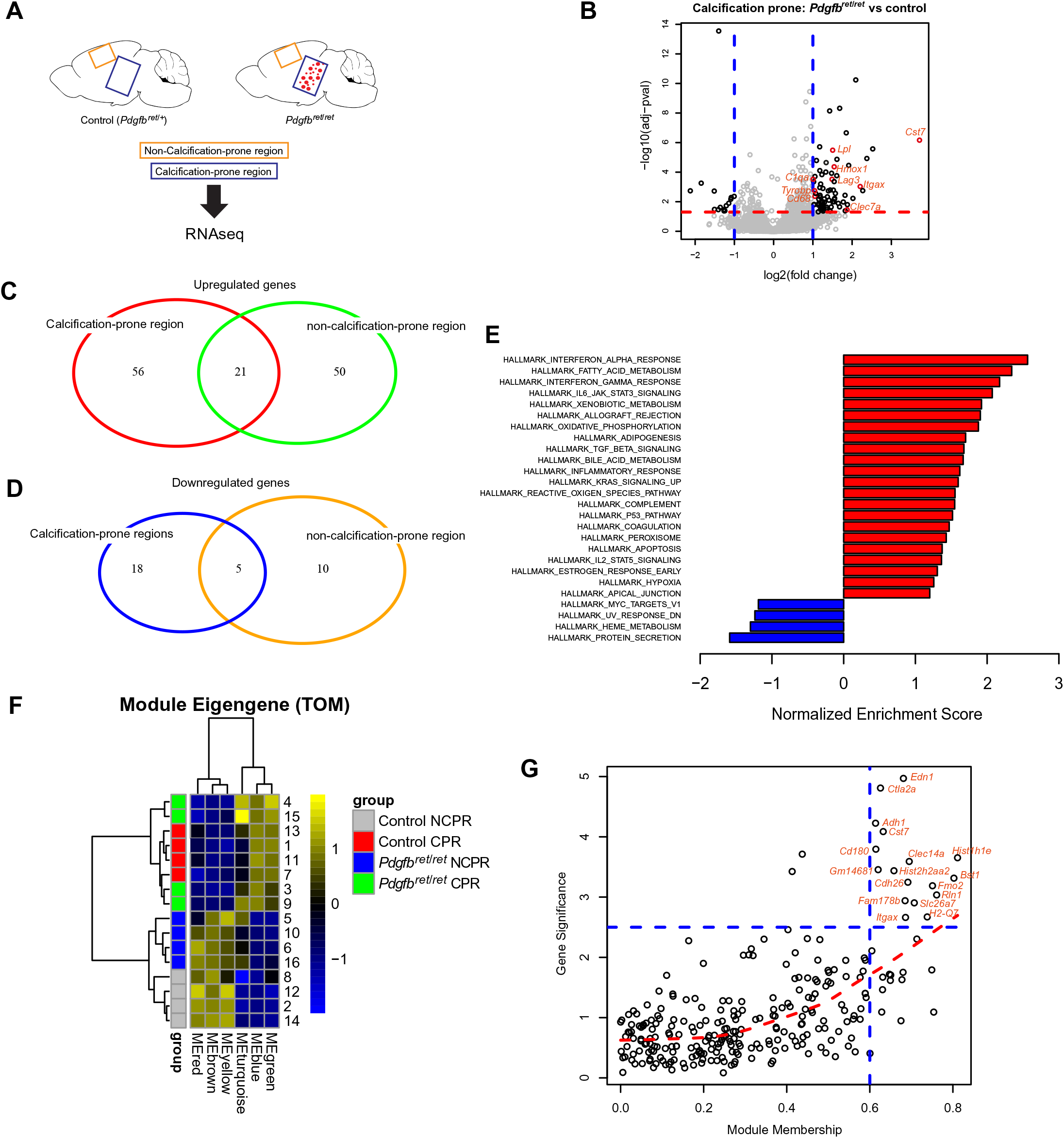
An inflammatory microenvironment surrounds vascular calcifications. **(A)** Samples collected for transcriptome analysis. RNA sequencing was performed from tissue isolated from two anatomical regions of *Pdgfb^ret/ret^* and control animals. Tissue enriched with brain calcifications was isolated from the thalamus/midbrain region labelled as “calcification-prone region”. Tissue isolated from the cortex is labelled “non-calcification prone region”. **(B)** Volcano plot showing deregulated genes in calcification-prone regions in *Pdgfb^ret/ret^* animals compared to control animals. **(C, D)** Venn diagrams showing upregulated **(C)** and downregulated **(D)** genes in calcification-prone and non-calcification prone regions. **(E)** Significantly upregulated (in red) and downregulated (in blue) pathways (p<0.05) in calcification-prone regions in *Pdgfb^ret/ret^* animals compared to controls. **(F)** For each module identified by the network analysis, the Module Eigengene (ME) was calculated, which summarizes the expression profile of the module. The turquoise module is associated with calcification-prone brain regions in *Pdgfb^ret/ret^* mice. CPR - calcification-prone region, NCPR – non-calcification-prone region **(G)** Graphical representation of hub genes with a high gene significance and module membership in the turquoise module.

**Figure 2.**
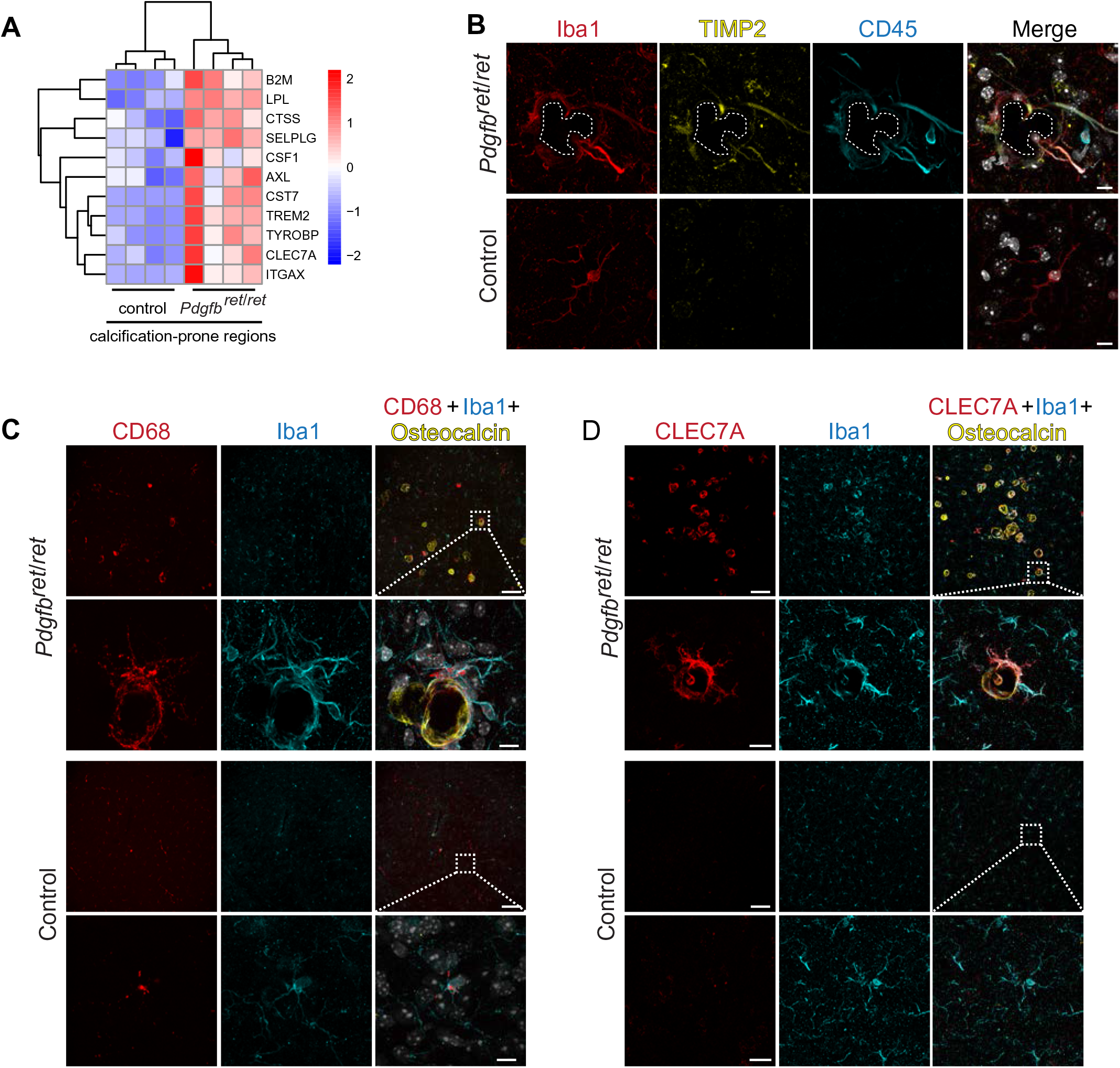
Microglia around vessel calcifications express proteins associated with a DAM/PAM signature. **(A)** Heatmap showing the expression of selected DAM signature genes significantly upregulated in calcification-prone regions (thalamus/midbrain) in *Pdgfb^ret/ret^* animals and controls. Data are presented as z-score. **(B-D)** Immunohistochemical validation of proteins associated with the DAM signature. **(B)** Iba1-positive (in red) microglia adjacent to a calcification (encircled with white dotted line) is strongly positive for CD45 (in cyan) and positive for TIMP2 (in yellow)**. (C)** Iba1-positive (in cyan) microglia surrounding osteocalcin positive calcifications (in yellow) are positive for CD68 (in red). **(D)** Iba1-positive (in cyan) microglia surrounding osteocalcin positive calcifications (in yellow) are positive for CLEC7A (in red). Nuclei were visualized using DAPI (in white). Scale bars – 10 μm (B, C inset, D inset) and 100 μm (C, D).

### Brain vessel-associated calcifications contain protein aggregates

Since CAM resemble amyloid β plaque-associated microglia in AD ^8^ (Fig. 2A-D) with protein deposits such as amyloid beta precursor protein (APP) and amyloid precursor-like protein (APLP2) ^42^, we next asked whether protein aggregation also occurs in brain calcifications. In order to investigate this possibility, we used Thioflavin T and Congo red, standard stains to detect amyloid fibrils in amyloid β plaques, and a recently developed amyloid interacting luminescent conjugated oligothiophene (LCO) - *h*-HTAA. Vessel-associated calcifications were Thioflavin T and Congo red (Suppl Fig. 2A) negative but positive for *h*-HTAA (Suppl Fig. 2B) that binds protein aggregates prior to the formation of amyloid fibrils recognised by Thioflavin T ^50^. Thus, vessel-associated calcifications contained aggregated proteins, which lack the β-pleated sheet conformation and structural regularity recognized by Thioflavin T or Congo red.

### Calcification-associated microglia (CAM) exhibit a distinct signature from DAM

Osteopontin (OPN, *Spp1*), one of the signature genes defining PAM and DAM ^8, 73, 74^, is deposited in brain calcifications in human PFBC and animal models of PFBC ^18^. Thus, we investigated if microglia surrounding calcifications express osteopontin and contribute to the previously reported deposition of osteopontin in brain calcifications ^18^. Immunohistochemical visualisation of microglia using anti-Iba1 and co-staining with osteopontin did not confirm osteopontin expression by microglia (Suppl Fig. 3A). Instead, osteopontin was expressed by a subset of GFAP-positive, reactive astrocytes surrounding or adjacent to calcifications (Suppl Fig. 3B). Hydroxyapatite, found in vessel-associated calcifications ^18^, and Aβ plaques induce formation of the inflammasome in macrophages and microglia, respectively ^75, 76^. We therefore investigated whether microglia around calcifications express ASC, an adaptor protein for inflammasome mediated caspase-1 activation. We could not find ASC positivity in microglia surrounding vessel calcifications in *Pdgfb^ret/ret^* animals (Suppl Fig. 3C), indicating that vessel-associated calcifications do not lead to inflammasome activation in microglia. Thus, CAM are distinct from DAM, although they share selected signature genes (Fig. 2A-D).

### Microglia proliferate around vessel-associated calcifications

In order to further characterize microglia in *Pdgfb^ret/ret^* mice, we performed flow cytometry analysis on calcification-prone brain regions using a multi-colour flow cytometry panel to distinguish immune cells and activated microglia from normal microglia. We detected eight immune cell clusters both in control and *Pdgfb^ret/ret^* mice as visualized by a dimension reduction algorithm Uniform Manifold Approximation and Projection (UMAP) ^77^ (Fig. 3A, Suppl Fig. 4A, B). Within the microglial cluster, we defined three sub-clusters (microglia, reactive microglia, and Ki67^+^ microglia) of microglia based on median marker expression of seven markers (CD11b, Ki67, F4/80, CD64, CX3CR1, CCR5, MerkTK) (Fig. 3B, C). Overall changes in *Pdgfb^ret/ret^* microglia compared to controls included increased expression of CX3CR1 and reduced CCR5 expression (Suppl Fig. 4C). We also detected an increase in Ki67^+^ microglia and the emergence of CX3CR1^high^, MerTK^high^ microglia (reactive microglia) in *Pdgfb^ret/ret^* animals (Fig. 3B, C). Using immunohistochemical staining for the proliferation marker Ki67, we confirmed the presence of proliferating CD45^high^ cells calcifications (Fig. 3D), while other cell types of the NVU (endothelial cells, pericytes, astrocytes) were Ki67 negative (Suppl Fig. 4D-F). Since Ki67 staining *ex vivo* can miss rarely proliferating cells, we performed *in vivo* labelling of proliferating cells using 5-ethynyl-2’-deoxyuridine (EdU) ^78^ in control and *Pdgfb^ret/ret^* mice. EdU positivity was only detected in microglial cells around calcifications (Fig. 3E). Thus, vessel calcifications evoke a proliferation of CAM, but not other cells at the NVU.

**Figure 3.**
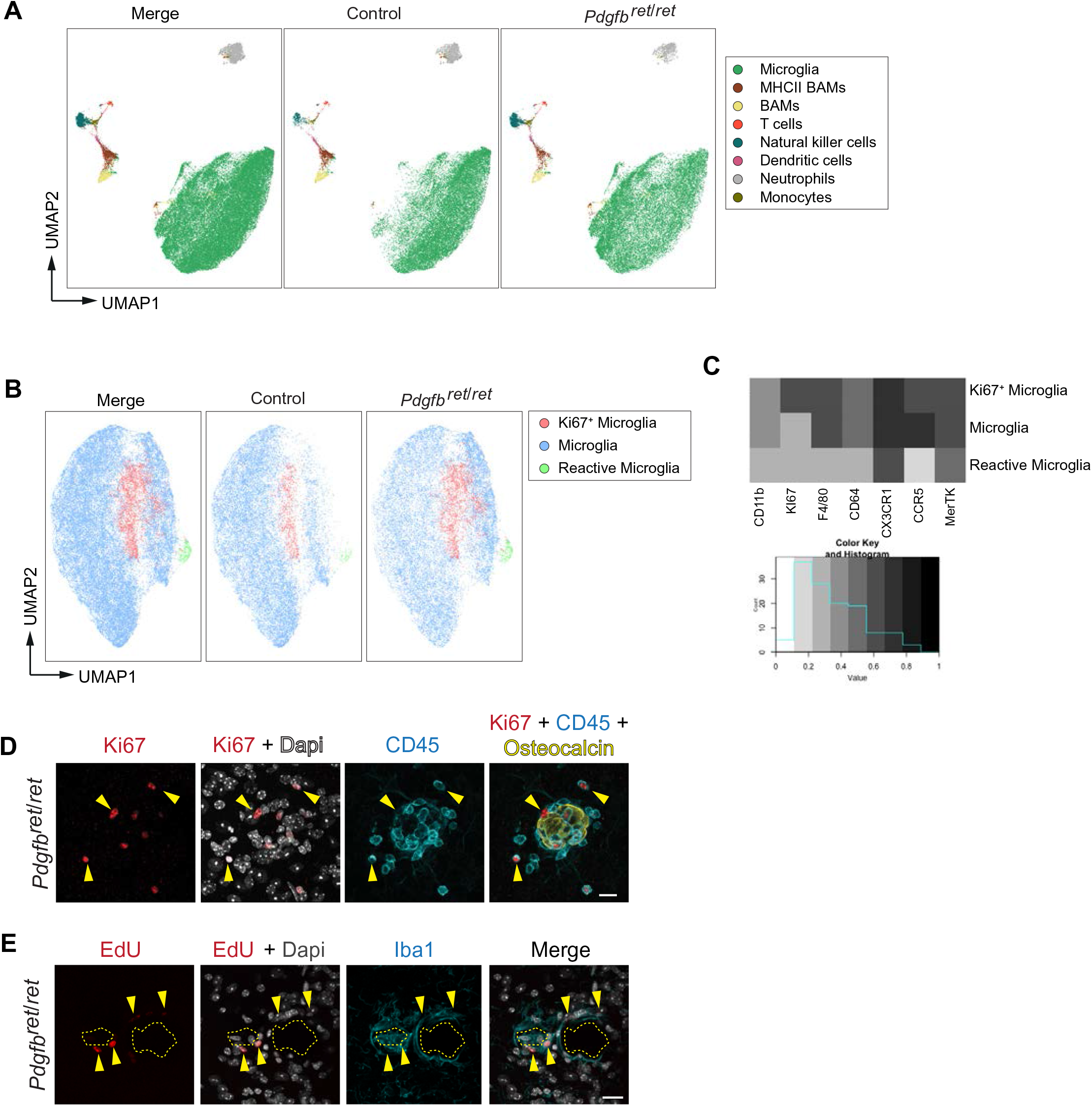
Microglia proliferate around vessel-associated calcifications. **(A-C)** Tissue from calcification-prone brain regions of *Pdgfb^ret/ret^* mice and controls were analysed using flow cytometry with a panel of antibodies to identify leukocytes and subtypes of brain myeloid cells. **(A)** Eight subpopulations of immune cells were identified with UMAP analysis. Microglia are depicted in green. **(B)** Microglia (in green in A) were isolated and analysed separately using UMAP. An increase in proliferating (Ki67^+^, in pink) microglia and the appearance of reactive (in green) microglia in *Pdgfb^ret/ret^* mice were observed in contrast to control animals (*Pdgfb^ret/wt^*). **(C)** Heatmap showing the expression of markers in different microglial sub-populations used to define sub-populations in B using UMAP. **(D)** A subset of CD45-positive cells (in cyan, yellow arrowheads) surrounding an osteocalcin positive brain calcification (in yellow) immunolabel with the proliferation marker Ki67 (in red). **(E)** EdU (in red) labelled Iba1-positive (in cyan, yellow arrowheads) microglia around brain calcifications (encircled with yellow dotted line). Nuclei were visualized using DAPI (in white, D and E). Scale bars – 15 μm (D), 10 μm (E).

### Microglia give rise to cathepsin K-expressing cells around vessel calcifications

We have reported that a subset of cells surrounding calcifications express osteoclast-associated markers - RANK and cathepsin K ^18^. In addition, cathepsin K, which is a principal collagen I degrading protease in bone, is deposited in brain calcifications ^18^. Here we show that RANK-expressing cells around calcifications are positive for the microglial marker Iba1 (Suppl Fig. 5). In order to investigate if resident microglia give rise to osteoclast-like cells around vessel-associated brain calcifications in *Pdgfb^ret/ret^* mice, we crossed two inducible Cre-lines (*Sall1-* CreER^T2^ and *Cx3cr1-*CreER^T2^) and a reporter line expressing tdTomato under the Rosa26 promoter (Ai14) with *Pdgfb^ret/ret^* animals to genetically label microglia. The *Cx3cr1-*CreER^T2^ line targets CX3CR1+ macrophages including microglia and “border-associated” macrophages in the CNS ^79^, whereas the *Sall1-*CreER^T2^ line targets resident microglia, but not infiltrating immune cells or brain perivascular macrophages ^45^. Mice were treated with tamoxifen at one month of age to induce tdTomato expression and sacrificed at four months of age (Fig. 4A). Immunostaining with cathepsin K and Iba1 showed a co-localization with tdTomato both when using *Cx3cr1-*CreER^T2^ (Fig. 4B) or *Sall1-*CreER^T2^ (Fig. 4C) lines, indicating that resident microglia respond to calcifications by expressing and depositing cathepsin K into vascular calcifications.

**Figure 4.**
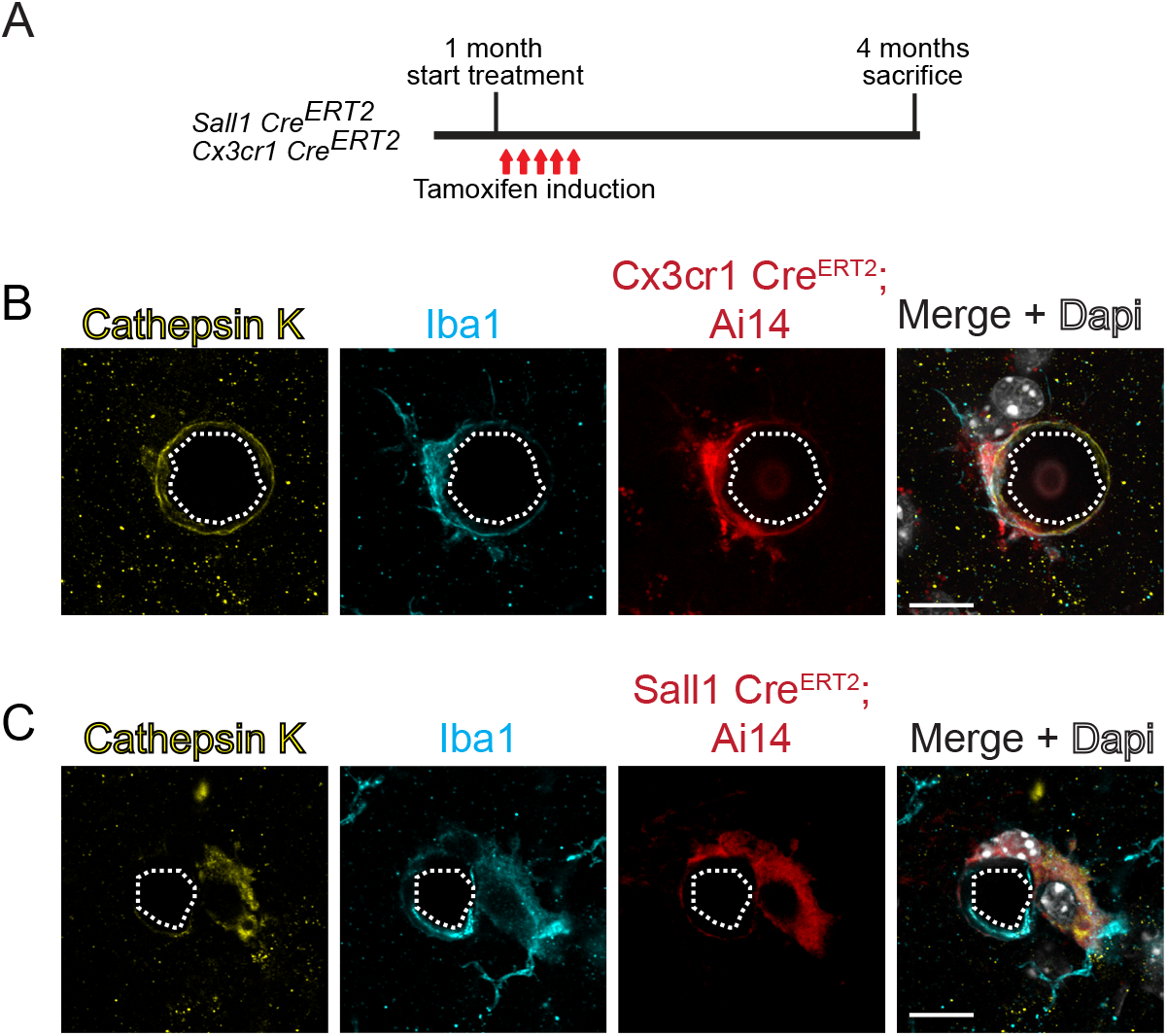
Microglia give rise to cathepsin K expressing cells around calcifications. **(A)** Experimental setup of lineage tracing experiments. At one-month of age control and *Pdgfb^ret/ret^* mice positive for *Sall1-*CreER^T2^; Ai14 or *Cx3cr1-*CreER^T2^; Ai14 were administered tamoxifen for 5 consecutive days and sacrificed at 4 months of age. TdTomato expression was induced using inducible Cre-lines where the expression of Cre is driven under *Sall1* or *Cx3cr1* promoter. **(B-C)** Cathepsin K staining (in yellow) co-localizes with tdTomato (in red) and Iba1staining (in cyan) adjacent to calcification (encircled with a white dotted line). Nuclei were visualized using Dapi (in white, in B and D). Scale bars – 10 μm (B, C).

### Pharmacological ablation of microglia intensifies vessel calcification

After establishing that cathepsin K-expressing cells surrounding vascular calcifications are derived from resident microglia (Fig. 4, Suppl Fig. 5), we asked if microglia actively participate in calcification-associated pathology. To this end, we depleted microglia in *Pdgfb^ret/ret^* and control animals for two months by using the colony stimulating factor 1 receptor (CSF1R) inhibitor, PLX5622 (Fig. 5A). In contrast to mice fed with control chow, mice fed with chow containing PLX5622 showed a reduction in the number of microglia (Suppl Fig. 6A, B). We used two markers, osteocalcin and APP, to detect and quantify vascular calcification (Fig. 5B). We previously reported that cerebral vascular calcifications are positive for bone matrix proteins, including osteocalcin ^18^. Moreover, a recent study identified that several non-bone proteins, including APP, are enriched in vessel-associated calcifications ^42^. A noticeable increase in vascular calcification (approx. 4 fold) in *Pdgfb^ret/ret^* mice was seen two months after microglial depletion (Fig. 5B, D). In addition, calcifications appeared heterogenous in size and shape in PLX5622-chow fed compared to control-chow fed *Pdgfb^ret/ret^* mice (Fig. 5B). In microglia-depleted *Pdgfb^ret/ret^* mice, the staining pattern of APP and osteocalcin was altered. APP and osteocalcin immunostaining was stronger along the periphery and weaker within calcifications. Some calcifications only stained for APP, whereas others immunolabeled only with osteocalcin (Fig, 5B, arrowheads). However, the ratio of APP and osteocalcin positive calcifications between PLX5622-treated and control chow-treated *Pdgfb^ret/ret^* mice was not statistically significant (Fig. 5C). CAM express cathepsin K (Fig. 4B, C). Accordingly, long-term microglial ablation should eliminate cathepsin K expression and deposition into calcifications. In fact, microglial depletion reduced cathepsin K deposition in calcifications (Fig. 5E, F). Consistent with our observation that activated microglia-encircling calcifications do not express osteopontin (Suppl Fig. 3), osteopontin is still deposited in calcifications in *Pdgfb^ret/ret^* mice after microglial depletion (Suppl Fig. 6C). Altogether, these results provide additional evidence that microglia are the principal cell type giving rise to osteoclast-like cells surrounding calcifications. In addition, our data show that microglial depletion aggravates the vessel-associated calcification phenotype, indicating that microglial activity could modify the pathophysiology of PFBC.

**Figure 5.**
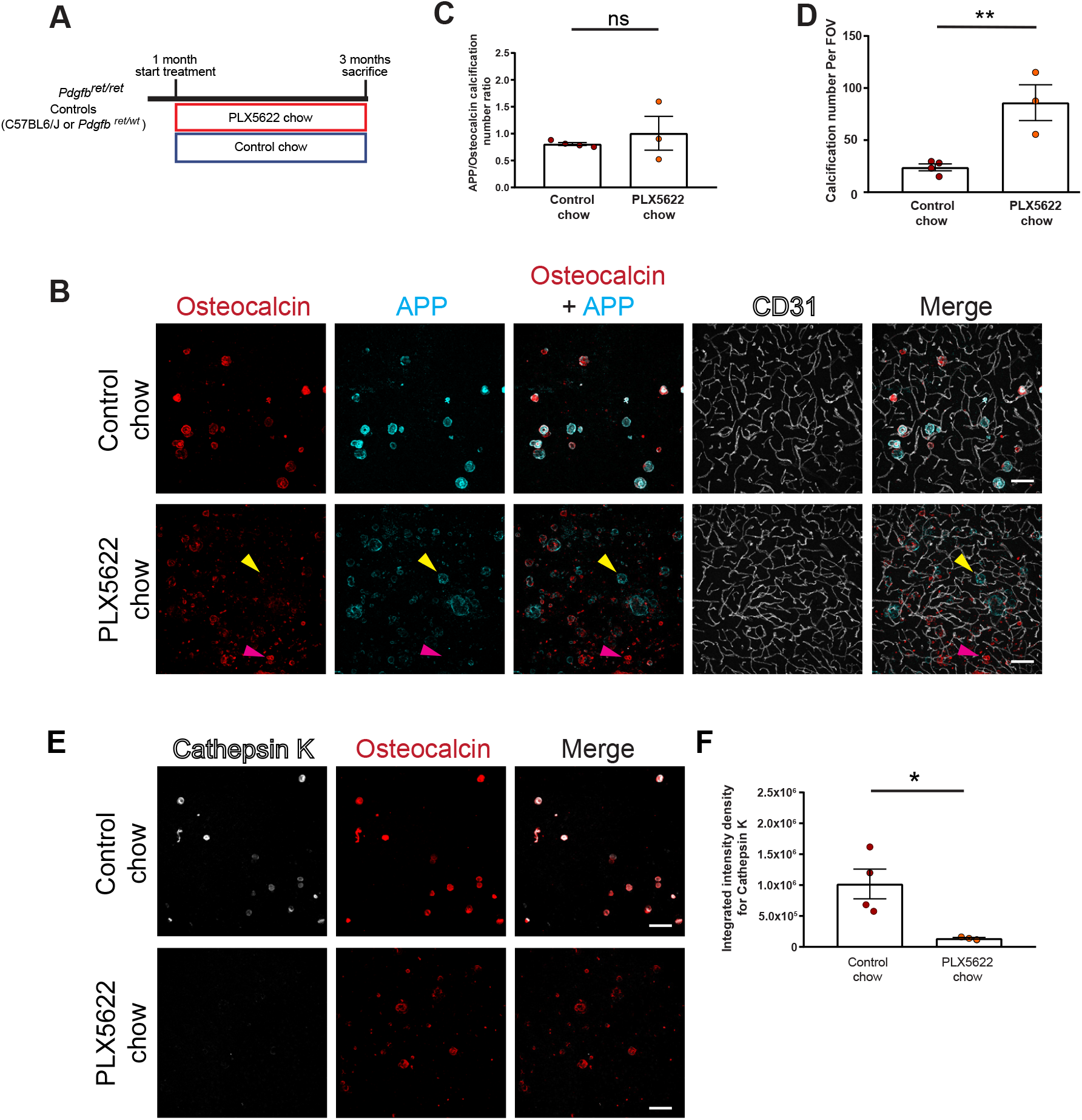
Pharmacological ablation of microglia aggravates vascular calcification. **(A)** Experimental setup of the pharmacological ablation of microglia. One-month old mice (*Pdgfb^ret/wt^*, *Pdgfb^ret/ret^* or C57BL6) were fed PLX5622 or control chow for two-months. Mice were sacrificed at 3 months of age. **(B)** Vessel-associated calcifications, visualised by osteocalcin (in red) and APP staining (in cyan), are increased in *Pdgfb^ret/ret^* compared to control chow fed *Pdgfb^ret/ret^* animals. Blood vessels are visualized using CD31 staining (in white). Note that some calcifications in PLX5622 treated *Pdgfb^ret/ret^* mice are only positive for APP (yellow arrowhead), whereas others are positive only for osteocalcin (magenta arrowhead). **(C)** The ratio between APP and osteocalcin positive calcifications after PLX5622 and control chow treatment. **(D)** Quantification of calcification number in *Pdgfb^ret/ret^* mice administered PLX5622 or control chow (unpaired two-tailed t-test, p = 0.0087). **(E)** Cathepsin K (in white) deposition in calcifications (in red) in *Pdgfb^ret/ret^* mice is reduced after PLX5622 treatment. **(F)** Quantification of cathepsin K intensity from immunohistochemical stains in (G; unpaired two-tailed t-test, p = 0.0276). Scale bars –100 μm (B, E). All data are mean ± SEM.

### Microglia ablation leads to bone protein containing axonal spheroids in white matter

After two months of chronic microglial depletion in control (C57BL6, *Pdgfb^ret/+^*) and *Pdgfb^ret/ret^* mice, (Fig. 5A), we observed conspicuous osteocalcin staining, indicative of calcification in the internal capsule, thalamus and striatum adjacent to white matter fiber tracts (Fig. 6A, dotted yellow area). However, linear osteocalcin-positive structures were not vessel-associated (Fig. 6B, Suppl Video 1). Negativity for alizarin red indicated that the observed white matter deposits were not calcified in three-month-old mice (Suppl Fig. 6D, upper pink panel). However, this result could be due the young age of the mice. We previously showed that vascular calcifications in *Pdgfb^ret/ret^* mice do not bind alizarin (i.e., are calcified) until four months of age ^22^. Accordingly, spheroidal deposits on vessels in the thalamus were alizarin red negative (Suppl Fig. 6D, mid blue panel, arrowheads), whereas occasional spheroids in the midbrain showed alizarin red positivity (Suppl Fig. 6D, lower orange panel). We further characterized deposits in white matter that appeared after microglial depletion using classical histochemical stains. Deposits were PAS positive, indicating the presence of glycoproteins (Fig. 6C, black arrowheads). The Bielschowsky silver stain revealed the presence of brown to black spheroids in white matter indicative of degenerating neurites (Fig. 6D, white arrowheads). In addition, co-staining of osteocalcin with APP, a marker for neuronal injury, and with myelin basic protein (MBP) showed co-localization within axonal spheroids (Fig. 6E, F). Furthermore, APP positive spheroids in white matter were also positive for another bone protein – osteopontin (Fig. 6G). Thus, microglial depletion leads to white matter injury, characterized by the formation of axonal spheroids, which immunolabel with APP and MBP as well as osteopontin, indicating the accumulation of bone proteins.

**Figure 6.**
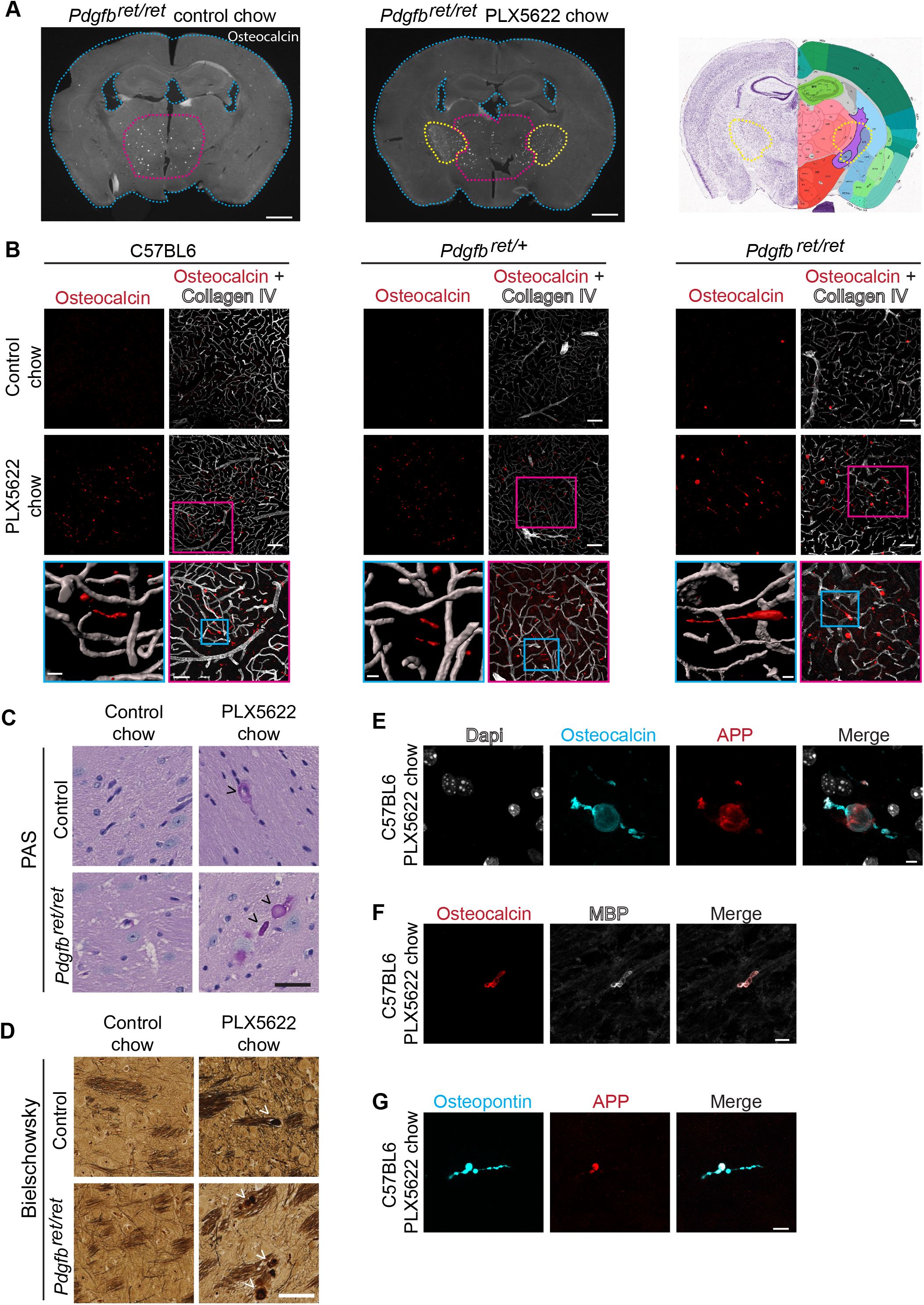
Microglial ablation leads to bone protein containing axonal spheroids in white matter. **(A)** Coronal sections of *Pdgfb^ret/ret^* mouse brains fed with control chow or chow containing PLX5622. Mice treated with PLX5622 exhibited conspicuous osteocalcin staining (in white) in the internal capsule (yellow dotted line, middle; black dotted line, right) in addition to calcifications in the thalamus (circled with a pink dotted line). Coronal mouse brain section depicts the location of analysed brain sections (Image credit: Allen Institute). **(B)** Linear inclusions, positive for osteocalcin (in red), in white matter are not vessel-associated in C57BL6 (left), *Pdgfb^ret/wt^* (middle) and *Pdgfb^ret/ret^* (right) mice. Blood vessels are visualized using the anti-collagen IV antibody (in white). **(C)** White matter deposits appearing after PLX5622 treatment *Pdgfb^ret/ret^* and control mice (*Pdgfb^ret/wt^*) are positive for PAS (black arrowheads). **(D)** Dystrophic neurites exhibiting axonal spheroids (indicated by white arrowheads) in white matter after PLX5622 treatment in *Pdgfb^ret/ret^* and control (*Pdgfb^ret/wt^*) mice. **(E-G)** White matter deposits appearing after microglial depletion are positive for osteocalcin (in cyan) and APP (in red) **(E)**, osteocalcin (in red) and MBP **(F)** (in white), and OPN (in cyan) and APP (in red) **(G)**. Nuclei were visualized using Dapi (E, in white). Scale bars – 1000 μm (A), 100 µm (B), 50 µm (B pink inset, C, D), 10 µm (B blue inset, F, G) and 5 µm (E).

### Microglia control vascular calcification in a TREM2-dependent manner

In addition to microglial depletion using a pharmacological CSF1R inhibitor, we used a genetic approach to impair microglial function in *Pdgfb^ret/ret^* mice by crossing *Pdgfb^ret/ret^* mice with *Trem2^-/-^* mice. Deletion of TREM2 leads to reduced microglial functionality ^80^. Furthermore, TREM2 expression is necessary for microglia to achieve the full DAM signature in a mouse model of AD ^8^. We noticed an altered pattern of calcification in *Pdgfb^ret/ret^;Trem2^-/-^* mice with vessels frequently encrusted with multiple osteocalcin-positive nodules (Fig. 7A, lower panels), yielding the ‘pearls-on-a-string’ phenotype reminiscent of human PFBC ^19^. This pattern was particularly evident in rostral thalamic regions that show less calcification than caudal thalamus and midbrain. Calcifications in *Pdgfb^ret/ret^;Trem2^+/+^* animals appeared as single nodules (Fig. 7A, upper panels) in accordance with the reported phenotype of calcifications in *Pdgfb^ret/ret^* mice ^18, 22, 42^. In addition to the altered pattern, vascular calcification was markedly aggravated in *Pdgfb^ret/ret^; Trem2^-/-^* and *Pdgfb^ret/ret^; Trem2^+/-^* mice (Fig. 7B). Similar to microglia depletion experiments, we observed altered APP and osteocalcin immunostaining of calcifications in *Pdgfb^ret/ret^* mice in the absence of one or two *Trem2* alleles (Fig. 7B). In *Pdgfb^ret/ret^; Trem2^+/+^* mice, calcifications stain uniformly with both antibodies used to visualize calcifications (osteocalcin and APP), whereas in *Pdgfb^ret/ret^; Trem2^+/-^* and *Pdgfb^ret/ret^; Trem2^-/-^* animals, calcifications showed weak APP or osteocalcin immunopositivity, respectively (Fig. 7B, D). Quantification of vascular calcifications (using both positivity for APP and OCN) in the midbrain of *Pdgfb^ret/ret^; Trem2^-/-^* and *Pdgfb^ret/ret^; Trem2^+/-^* mice showed an increased calcification (4-6 fold) compared to *Pdgfb^ret/ret^;Trem2^+/+^* mice (Fig. 7D). Thus, functional TREM2 in microglia is necessary to limit the formation of vessel calcifications in *Pdgfb^ret/ret^* mice. The lack of *Trem2* in control mice does not lead to brain calcification (Suppl Fig. 7A). Additionally, we observed that microglia surrounding calcifications express CLEC7A, a DAM-associated protein dependent on *Trem2*, in *Pdgfb^ret/ret^;Trem2^-/-^* animals (Suppl Fig. 7B). Interestingly, cathepsin K deposition into calcifications is TREM2-dependent, *Pdgfb^ret/ret^; Trem2^+/-^* and *Pdgfb^ret/ret^; Trem2^-/-^* mice displayed a strongly reduced cathepsin K deposition into calcifications (Fig. 7E, F). In summary, these data show that cathepsin K expression by CAM is TREM2-dependent and further corroborate that microglia control the growth of vascular calcifications in brain.

**Figure 7.**
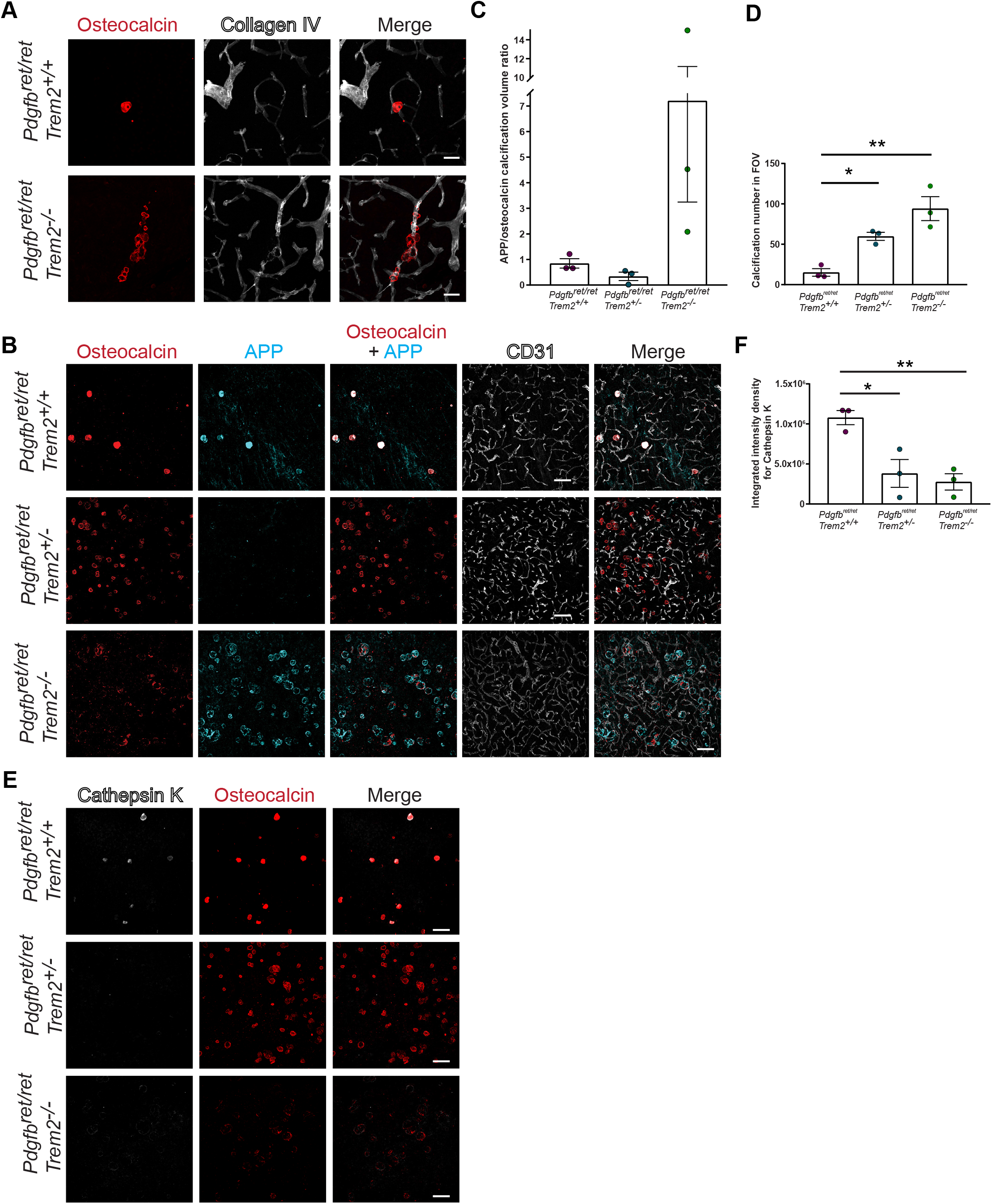
TREM2-dependent microglial function is crucial in controlling vascular calcification. **(A)** Altered pattern of vessel calcification in *Pdgfb^ret/ret^; Trem2^-/-^* animals compared to *Pdgfb^ret/ret^; Trem2^+/+^* animals. Calcifications are visualized using osteocalcin staining (in red) and blood vessels using collagen IV staining (in white). **(B)** *Pdgfb^ret/ret^; Trem2^-/-^* and *Pdgfb^ret/ret^; Trem2^+/-^* animals show an increase in calcification load when compared to *Pdgfb^ret/ret^; Trem2^+/+^* animals. Vascular calcification is visualized with osteocalcin (in red) and APP (in cyan) staining, and blood vessels using CD31 staining (in white). Note the staining of the two selected calcification markers in *Pdgfb^ret/ret^* animals that differ in zygosity for *Trem2*. **(D)** Ratio between APP and osteocalcin staining. **(C)** Quantification of calcification using immunohistochemical stains (one-way ANOVA with Dunnett’s multiple comparison; *p = 0.0270, **p = 0.0018). **(E)** Cathepsin K (in white) deposition in calcifications (in red) is reduced in *Pdgfb^ret/ret^; Trem2^-/-^* and *Pdgfb^ret/ret^; Trem2^+/-^* animals when compared to *Pdgfb^ret/ret^; Trem2^+/+^* animals. **(F)** Quantification of cathepsin K intensity with immunohistochemical stains (one-way ANOVA with Dunnett’s multiple comparison; *p = 0.0146, **p = 0.0076). Scale bars – 50 μm (A) and 100 μm (B, E). All data are means ± SEM.

### Microglia depletion or functional modulation alters astrocyte reactivity but not the neurotoxic profile around calcifications

We next explored whether microglia also modify the strong astrocyte reactivity around calcifications ^18, 22^. We had shown previously that astrocytes encircling brain calcifications exhibit a neurotoxic response (e.g., C3, LCN2 expression) as well as an unusual reactive phenotype (e.g., podoplanin expression) ^18^. We investigated if astrocyte reactivity and the expression of neurotoxic markers is altered after modifying the number and function of microglia in *Pdgfb^ret/ret^* mice. Microglia ablation using PLX5622 or compromised function (*Trem2* genotype) in *Pdgfb^ret/ret^* mice resulted in an altered staining pattern of GFAP and podoplanin, proteins expressed by reactive astrocytes surrounding calcifications (Fig. 8A, B). In *Pdgfb^ret/ret^* mice treated with PLX5622, GFAP reactivity showed a diffuse pattern, most likely because of an increased density of calcifications, compared to mice that received control chow (Fig. 8A, E, G). Furthermore, similar alterations in GFAP staining were observed in *Pdgfb^ret/ret^* mice with impaired microglial function: *Pdgfb^ret/ret^; Trem2^+/-^* and *Pdgfb^ret/ret^; Trem2^-/-^* (Fig. 8B). Podoplanin staining in reactive astrocytes surrounding calcifications was markedly reduced (Fig. 8A, B). We quantified podoplanin and GFAP staining intensity and calculated the staining intensity ratio, which showed a trend towards a reduction in *Pdgfb^ret/ret^* mice treated with PLX5622 (Fig. 8C) and a significant reduction in *Pdgfb^ret/ret^; Trem2^+/-^* and *Pdgfb^ret/ret^; Trem2^-/-^* mice (Fig. 8D) compared to *Pdgfb^ret/ret^* mice. Notably, we did not detect a reduction in the expression of neurotoxic signature markers – LCN2 and C3 by reactive astrocytes surrounding calcifications after microglia depletion in *Pdgfb^ret/ret^* mice (Fig. 8E, F). We quantified LCN2 and C3, and GFAP staining intensity and calculated the staining intensity ratio, which showed no significant difference between PLX5622-treated and non-treated *Pdgfb^ret/ret^* mice (Fig. 8G, H). Altogether, these results indicate that microglia modulate astrocyte reactivity but not required to evoke a neurotoxic astrocyte phenotype in response to vessel calcification.

**Figure 8.**
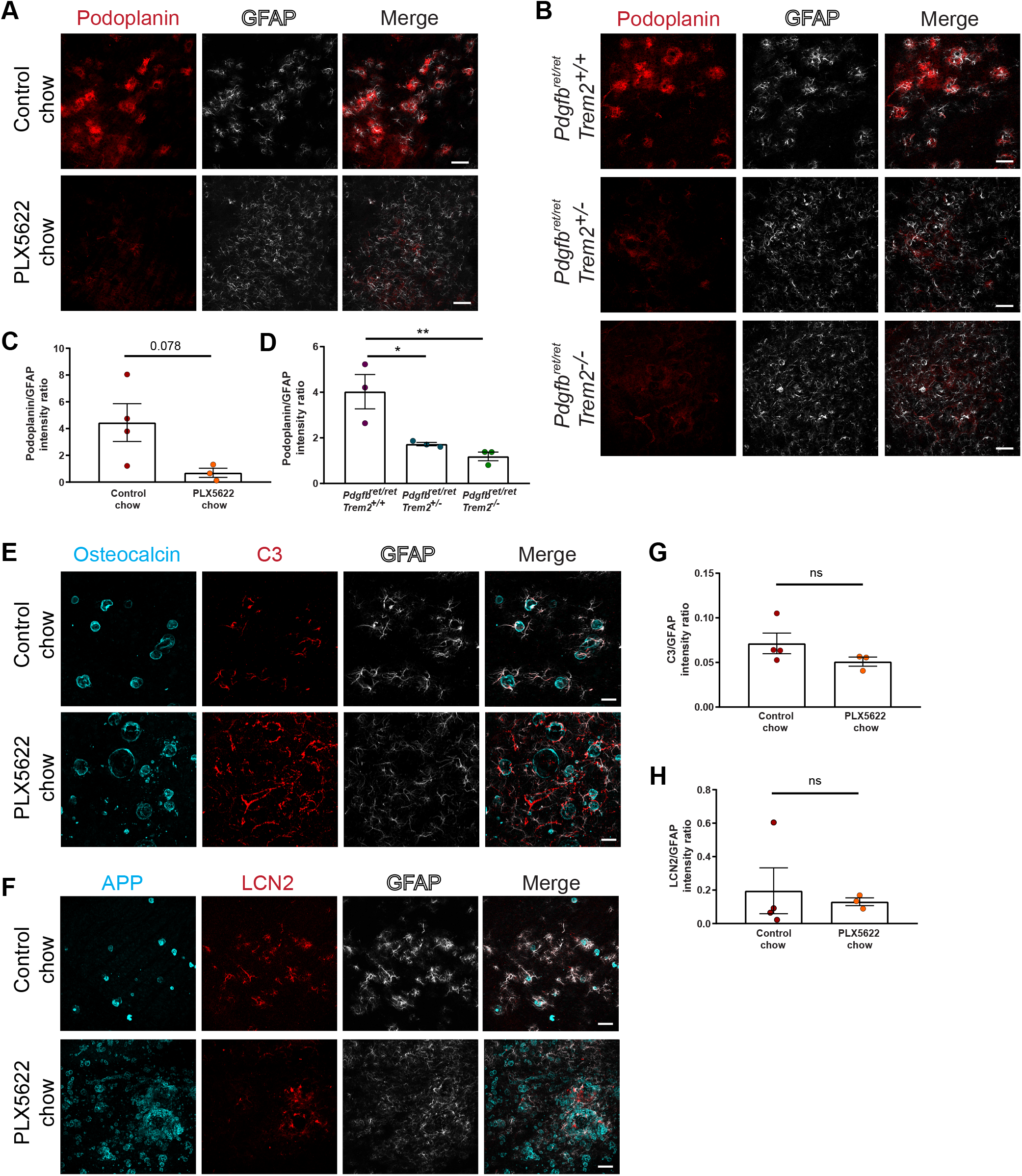
Alterations in microglia number and function change astrocyte reactivity. **(A, B)** Podoplanin (in red) expression in GFAP-positive astrocytes surrounding calcifications (in cyan) is reduced in *Pdgfb^ret/ret^* animals treated with PLX5622 **(A)** or lacking *Trem2* **(B)**. **(C, D)** Quantification of podoplanin intensity normalized to GFAP intensity in mice treated with PLX5622 (unpaired two-tailed t-test, p=0.078) **(C)** and with different zygosity for *Trem2* (one-way ANOVA with Dunnett’s multiple comparison; *p=0.008, **p=0.02) **(D)**. **(E)** C3 (in red) expression in GFAP-positive astrocytes (in white) around calcifications (osteocalcin, in cyan) in *Pdgfb^ret/ret^* animals treated with PLX5622. **(F)** LCN2 (in red) expression in GFAP-positive astrocytes (in white) around calcifications (APP, in cyan) in *Pdgfb^ret/ret^* animals treated with PLX5622. **(G)** Quantification of C3 expression in astrocytes (unpaired two-tailed t-test, p=0.21). **(H)** Quantification of the LCN2 expression in astrocytes (Mann-Whitney two-tailed test p=0.63). Scale bars – 100 μm (A, C) and 50 μm (E, G). All data are means ± SEM.

## Discussion

Our results demonstrate that in addition to already known microglial functions, microglial activity in the context of vascular calcification is beneficial by identifying incipient harmful calcification (Fig. 9). We have used a pharmacological approach in a mouse model for PFBC to modify microglial numbers and a genetic approach to modify microglial function. Both approaches enhance calcification of the NVU (Fig. 5, 7). Vascular calcification, resulting in increased pulse wave velocity and decreased end-organ perfusion and damage, is very common with aging as a major risk factor ^81^. Surprisingly, studies investigating vascular calcifications in the brain are rare ^82^. Therefore, the consequences of calcification on vascular function at the NVU, including the neurovascular coupling, have not been studied. As noted, brain calcification is a common incidental CT finding ^28, 29^. It is plausible that under homeostatic conditions, i.e., without underlying vascular pathology, microglia limit vascular calcification during aging (Fig. 9), which should be addressed by future studies. Vascular calcification is commonly observed in neurodegenerative diseases such as AD ^83^, which warrants further investigation to determine whether microglial dysfunction contributes to vascular pathology in AD. Thus, in addition to parenchymal surveillance and the removal of various injurious stimuli, microglia could remove calcium phosphate precipitates and halt calcification of the NVU (Fig. 9).

**Figure 9.**
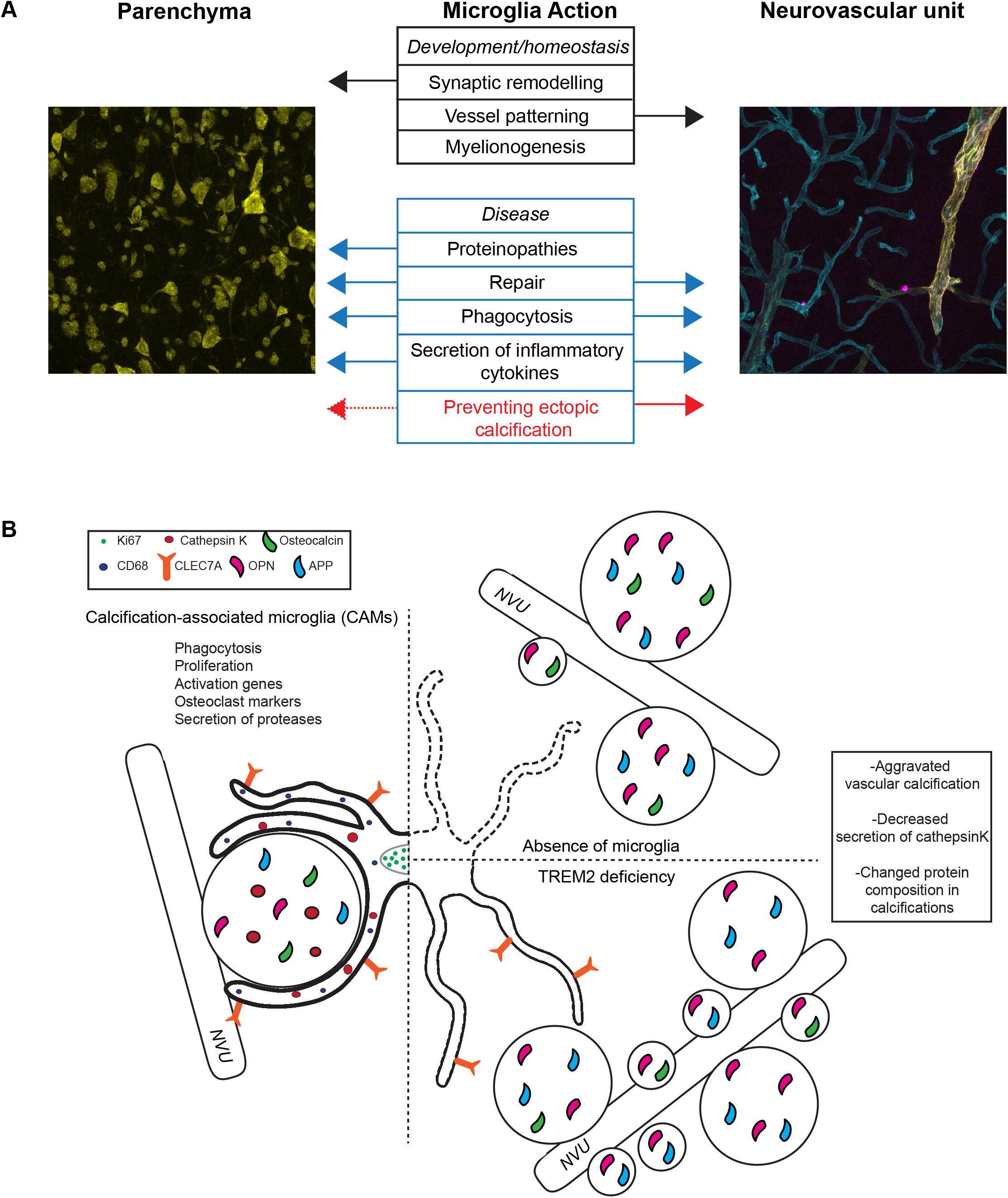
Microglia functions in brain parenchyma and at the NVU. **(A)** Microglia function during development, homeostasis and in disease states with their respective action on parenchymal and NVU cells. Text in red marks a new function identified in this study. **(B)** Calcified protein aggregates at the NVU are surrounded by calcification-associated microglia (CAM). CAMs are phagocytotic (express CD68; blue circles), proliferative (express Ki67; green circles), activated (express CLEC7A; orange receptor), and secrete ECM degrading proteases (cathepsin K, red circles). Vascular calcifications contain bone proteins (osteocalcin, green teardrop; osteopontin, pink teardrop) and non-bone proteins (APP, light blue teardrop; cathepsin K, red circles). Microglial depletion (indicated by a dotted line) or genetic deletion of *Trem2* in *Pdgfb^ret/ret^* mice leads to increased calcification of the NVU and changes in the protein composition (e.g., lack cathepsin K).

Microglial activation is insult-dependent ^70^; however, some microglial activation gene signatures are shared (e.g. *Spp1, Clec7a*) by several neurodegenerative proteinopathies, developmental stages and aging-related changes ^8, 10, 66, 70^. We observed that CAMs express CLEC7A (Fig. 2 D), but not osteopontin (OPN, gene name: *Spp1*) (Suppl Fig. 3A) indicating that calcifications elicit a microglial response similar to but distinct from DAM/PAM and aging microglia. CLEC7A is still expressed by CAMs in the absence of *Trem2* (Suppl Fig. 7B), which is necessary for the induction of the DAM signature in microglia ^8^. Furthermore, osteopontin is deposited in vessel calcifications after microglial depletion (Suppl Fig. 6C). We detected osteopontin expression in a subset of GFAP-positive astrocytes surrounding calcifications (Suppl Fig. 3B), similar to osteopontin expression in reactive astrocytes after a stab wound injury ^84^. Osteopontin is highly expressed in injured tissues, including brain diseases (e.g., Parkinson disease, multiple sclerosis) ^85–89^. One of the biological functions of osteopontin is to act as a hydroxyapatite binding mineral chaperone to inhibit the formation of hydroxyapatite crystals ^90^. The induction of osteopontin by different cells (e.g. microglia, astrocytes, neurons) in response to various brain insults ^84, 87, 91, 92^ could prevent injury and ectopic calcification in the parenchyma or at the NVU. Ectopic soft-tissue calcifications in the periphery and in brain, which are induced by excitotoxic insults, have a similar lamellar structure with darker appearing lamellae rich in osteopontin ^86, 93^. Notably, *Spp1^-/-^* mice develop severe secondary neurodegeneration accompanied by increased brain calcification in response to excitotoxic insults ^94^.

We show that cathepsin K deposition in vascular calcifications is decreased after altering microglial numbers or function in *Pdgfb^ret/ret^* mice (Fig. 5E and 7E). Accordingly, cathepsin K-expressing cells surrounding vessel calcifications derive from resident microglia (Fig. 4B, C). In bone, cathepsin K is expressed and secreted by osteoclasts and represents the primary enzyme for degrading type I collagen ^95^. Our work and that of others have documented collagen I in brain calcifications ^18, 96^. Thus, cathepsin K activity could be necessary for degrading the extracellular matrix (ECM) required for calcium phosphate precipitation. Microglia-specific modification of cathepsin K expression should clarify the role of cathepsin K in vascular calcification. Microglial activity that impedes vessel calcification could trigger a coordinated activation of several pathways in protein and hydroxyapatite degradation. However, it is currently unclear if microglia remove hydroxyapatite present in vascular calcifications ^18^. Macrophages have been shown to remove various crystals, including hydroxyapatite, which leads to activation of the NLRP3 inflammasome pathway ^97^. Our studies show that although vessel calcifications elicit strong microglial reactivity *in vivo,* they do not activate the inflammasome (Suppl Fig. 3C). Further studies are needed to understand how microglia sense and remove vascular calcification and whether this activates intracellular pathways as those evoked by self-derived damage-associated molecular patterns (DAMPs) via pattern recognition receptors. In addition to altered cathepsin K deposition, we observed an altered deposition of APP and osteocalcin into calcifications in *Pdgfb^ret/ret^* crossed with *Trem2^-/-^* or *Trem2^+/-^* or after microglia depletion (Fig. 5B, 7B). This finding indicates that microglial activity not only impedes the growth of calcifications but also modifies matrix composition of calcifications.

In this study, we observed that functional TREM2 is required to halt vessel-calcification in a mouse model of PFBC (Fig. 7). TREM2 deficiency also leads to an altered vessel calcification pattern in *Pdgfb^ret/ret^* mice (Fig. 7A), similar to that described in human autopsy cases. Vascular calcification in PFBC is sometimes described as ‘pearls on a string’ due to numerous, tiny spherical calcifications that encrust almost the entire abluminal side of capillaries ^19, 98^. Interestingly, calcifications appear as single nodules ^18, 20, 22, 24, 42^ in mouse models of PFBC. Variants in a microglia-specific gene, *TREM2*, have been associated with an increased risk for AD ^99^. Studies in mouse models of AD have linked functional TREM2 to the development of a specific microglia activation state - DAM ^8, 9^, and to maintaining microglial metabolic fitness as well as a sustained microglial response to Aβ-plaque-induced pathology ^100^. It has been proposed that even though microglia are efficient in phagocytosing protein aggregates in the early stages of AD, over time they lose their efficiency ^101^. Therefore, it is plausible that by crossing *Pdgfb^ret/ret^* with *Trem2^+/-^* or *Trem2^-/-^*, vascular calcification is accelerated due to microglial exhaustion ^101^. It is noteworthy that microglia encircling calcifications contain numerous phagocytotic vesicles ^42^. Further studies are needed to understand whether TREM2-driven microglial function is initiated due to altered proteostasis and calcification at the NVU or neuronal death occurring due to altered vessel function. These processes, however, are not necessarily mutually exclusive.

Several lines of evidence indicate that cell-autonomous defects in microglia lead to brain disease and cerebral calcification ^31, 32, 37, 38, 40^. Microglia have also emerged as a disease modifier in a wide range of neurodegenerative diseases (e.g. AD, FTD) ^99, 101, 102^. Of five genes implicated in PFBC (*PDGFB, PDGFRB, SLC20A2, XPR1, MYORG*), microglia express PDGFB, SLC20A2, XPR1 ^10, 66^. Although microglia express PDGFB, they do not express the receptor – PDGFRB. Thus, it is unlikely that *PDGFB* or *PDGFRB* haploinsufficiency in PFBC causes cell-autonomous microglial dysfunction. Phosphate importer SLC20A2 and exporter XPR1 are expressed ubiquitously in the brain ^103–106^ and their specific role in microglia has not been investigated. However, it has been reported that in zebrafish - *xpr1b*, an orthologue of *XPR1,* is crucial for the differentiation of tissue-resident macrophages and microglia ^107^. In mice, the full knockout of *Xpr1* is embryonic lethal ^108^, but the embryonic phenotype has not been characterized. Thus, further studies are needed to dissect the role of PFBC genes in microglial function.

In this study, based on positivity for osteocalcin, we found by serendipity that microglial reduction by chronic CSF1R inhibition using PLX5622 for two months resulted in localized axonal damage to fiber tracts of the internal capsule and adjacent thalamic and striatal areas (Fig. 6). Positive staining for osteocalcin and osteopontin coincided with the presence of dystrophic neurites exhibiting spheroid formation (Fig. 6E-G), similar to the pathology described in patients with leukoencephalopathy caused by *CSF1R* mutations ^109^. Interestingly, these patients exhibit brain calcification in white matter regions ^40^ as well as a reduction in microglia in affected regions ^110^. It is plausible that axonal spheroids become calcified during the course of the disease. Further studies are needed to understand the relationship between the appearance of axonal spheroids positive for bone proteins and white matter calcification. We observed that inclusions within dystrophic neurites stain positive for MBP (Fig. 6G), a protein secreted by oligodendrocytes, indicating a disrupted homeostasis in oligodendrocytes. Previous studies on white matter microglia have shown that microglia promote myelinogenesis during early development, providing evidence for a role in optimizing oligodendrocyte function ^111, 112^. Thus, further studies are needed to ascertain whether axonal damage is directly caused by a reduction in microglia or the absence of microglia has an effect on another cell type (e.g. oligodendrocytes), which may enhance spheroid formation in neurites.

In conclusion, we describe a novel, unrecognized role of microglia in brain vascular calcification. In addition, we show that functional microglia are important to prevent calcification of the NVU in neurodegenerative diseases with a compromised NVU. Proposed mechanisms by which microglia control vascular calcification include the removal of apoptotic cells and spontaneous calcium-phosphate precipitates as well as the prevention of nucleation of hydroxyapatite by controlling proteostasis of the ECM, and/or the secretion of anti-calcifying proteins or molecules.

## Acknowledgements

This work was supported by the Swiss National Science Foundation (grants 31003A_159514 to AK, PP00P3_170626 and BSGI0_155832 to MG), the Synapsis Foundation, Fonds zur Förderung des akademischen Nachwuchses (Zurich University), the Leducq Foundation (SphingoNet), the Swiss Heart Foundation, The Swiss Cancer League to AK, and Forschungskredit und Stiftung für Forschung an der Medizinischen Fakultät der Universität Zürich (grant no. FK-16-034) to YZ. Imaging was performed with equipment maintained by the Center for Microscopy and Image Analysis, University of Zurich and RNA sequencing was performed at the Functional Genomics Center Zurich, University of Zurich and ETH. We thank Plexxikon for providing PLX5622, F. Franzoso for technical help and S. Hornemann for discussions.

