## Supplementary Figures for "Microglia control small vessel calcification via TREM2"

### Supplementary Figure 1.

Zarb et al.

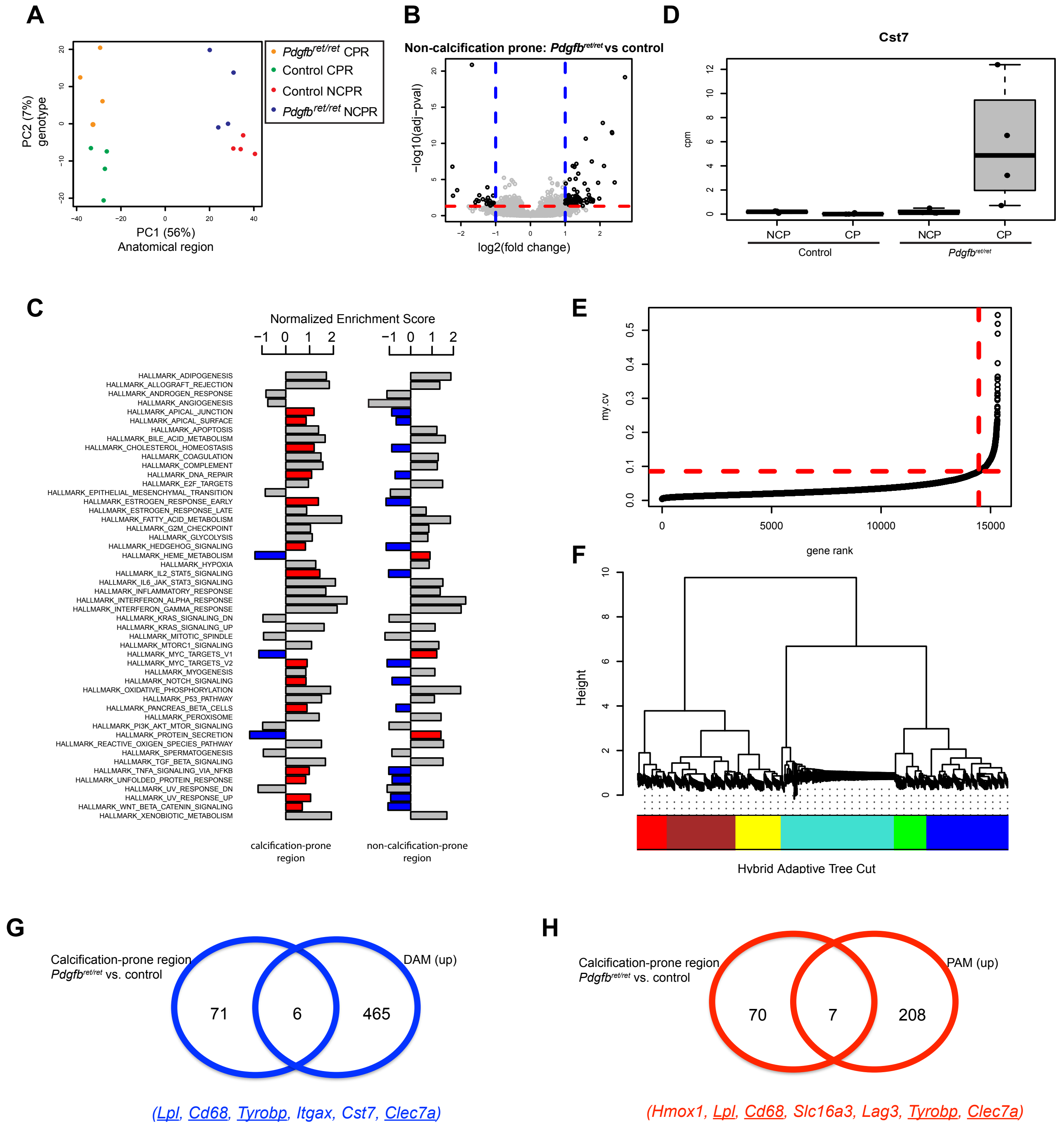

**Supplementary figure 1. Transcriptomic analysis of brain calcifications.**

**(A)** Principle component (PC) analysis of the samples after batch adjustment. Most of the data set variation (PC1, 56 % of variation) is due anatomical location (non-calcification prone regions (NCPR), cortex) vs. calcification-prone regions (CPR), thalamus/midbrain) and sample genotype, which accounts for the 7% of variation (PC2). **(B)** Volcano plot showing deregulated genes within the cortex (non-calcified region) in *Pdgfb*$^{ret/ret}$ and control animals. **(C)** Signalling pathways in *Pdgfb*$^{ret/ret}$ mice that are inversely enriched in a calcification-prone region (thalamus) compared to a non-calcification prone region (cortex). **(D)** Boxplot showing the *Cst7* expression in different samples. **(E)** Selection criteria for the network analysis of highly variable genes. **(F)** Co-expression networks were generated with 867 highly variable genes (selected in **E**). Hierarchical clustering of the topological overlap matrix dissimilarity further revealed 6 modules of positively correlated genes; each was assigned a colour as a label. **(G, H)** Venn diagram showing the overlap between upregulated genes (FDR <0.05 and fold change >2) in calcification-prone regions in *Pgfb*$^{ret/ret}$ mice and disease-associated microglia (DAM) **(G)** or proliferative-region associated microglia (PAM) **(H)**. Underlined genes are shared between PAM and DAM signatures.

### Supplementary Figure 2.

Zarb et al.

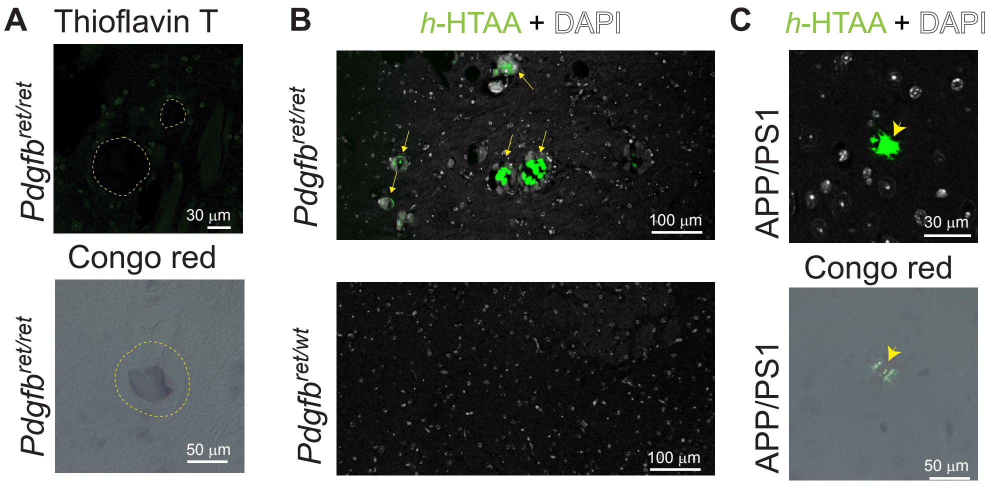

**Supplementary figure 2. Brain vessel-associated calcifications contain protein aggregates** Brain calcifications are negative for Thioflavin T and Congo red **(A)**, but positive for *h*-HTAA staining (yellow arrows). **(B)**. Brain sections from 8 months old APP/PS1 mice used as a positive control for *h*-HTAA and Congo red stainings (yellow arrowhead).

### Supplementary Figure 3.

Zarb et al.

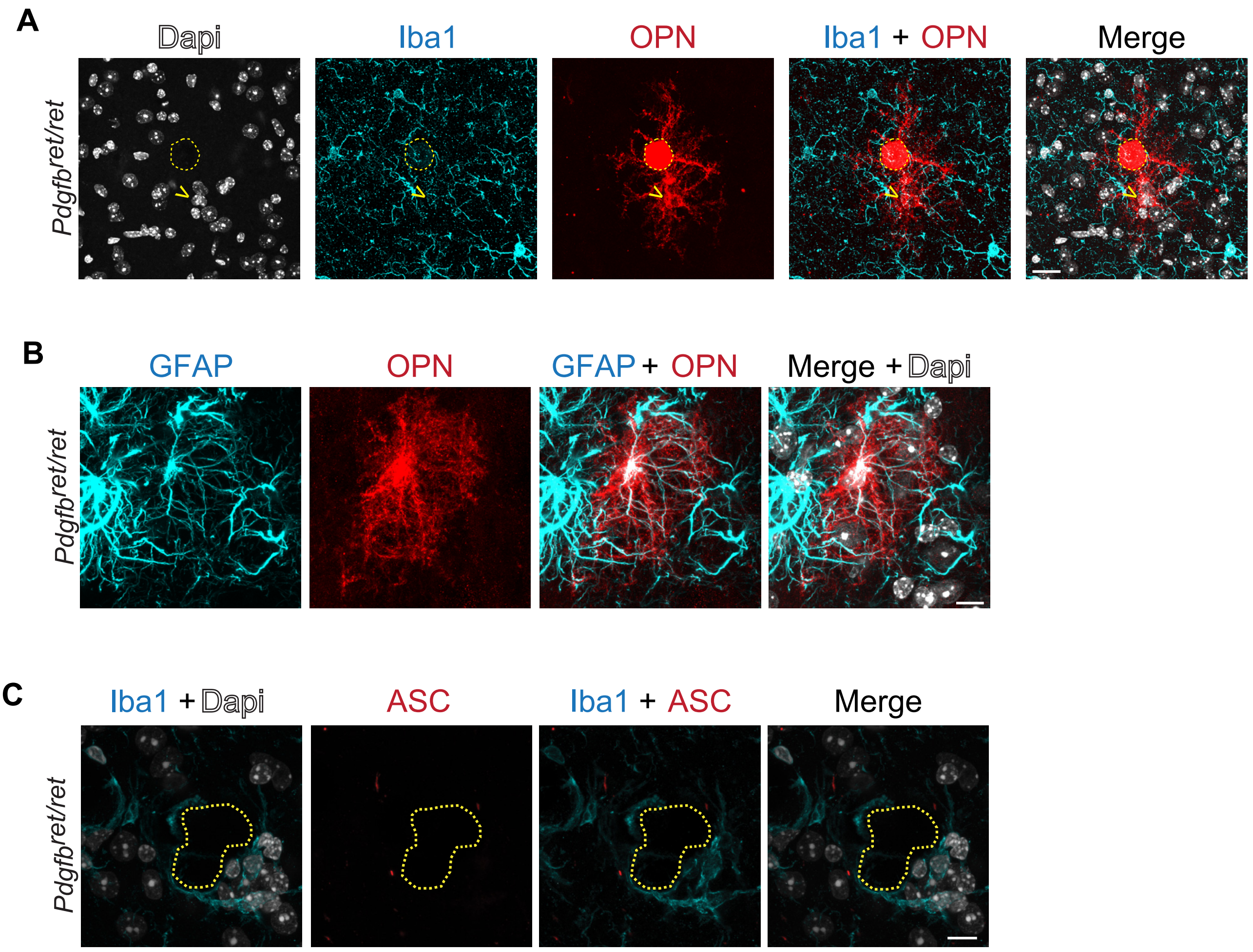

**Supplementary figure 3. Calcification-associated microglia do not express osteopontin or show inflammasome activation.**

**(A)** Microglia (Iba1, in cyan) around a brain calcification (encircled with yellow dotted line) are not positive for osteopontin (OPN) (in red). Osteopontin-positive cell is indicated with a yellow arrowhead. **(B)** Astrocytes (GFAP, in cyan) in the thalamus of *Pdgfb*$^{ret/ret}$ mice express osteopontin (OPN, in red). **(C)** Iba1-positive (in cyan) microglia surrounding brain calcifications (encircled with yellow dotted line) do not express ASC (in red), a component of the inflammasome. Nuclei were visualized using Dapi (in white). Scale bar – 20 µm (A), 10 µm (B, C).

### Supplementary Figure 4.

Zarb et al.

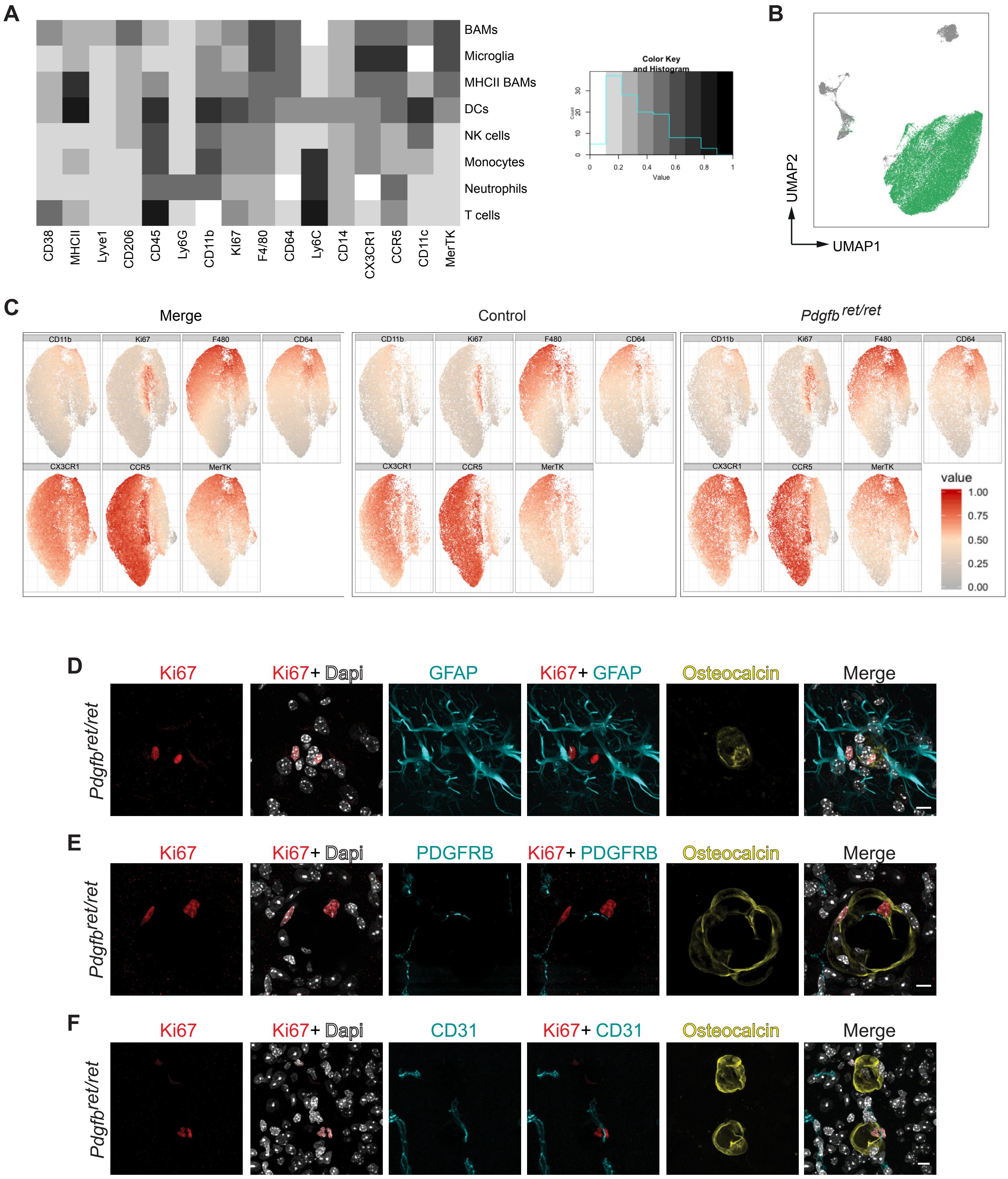

**Supplementary figure 4. Microglia reactivity and identification of Ki67$^{+}$ cells surrounding brain calcifications in *Pdgfb*$^{ret/ret}$ animals.**

**(A)** Heatmap showing mean marker expression of markers to identify leukocyte populations by flow cytometry analysis. **(B)** UMAP plot of leukocyte populations detected in calcification-prone regions of *Pdfgb*$^{ret/ret}$ and control animals. Microglial cluster is in green. **(C)** Median marker expression within the microglial cluster. **(D-F)** Ki67-positive (in red) cells around brain calcifications (osteocalcin, in yellow) are not **(D)** astrocytes (GFAP, in cyan), **(E)** mural cells (PDGFRB, in cyan) **(F)** or endothelial cells (CD31, in cyan). Nuclei were visualized using DAPI (in white). Scale bar - 10 µm.

### Supplementary Figure 5.

Zarb et al.

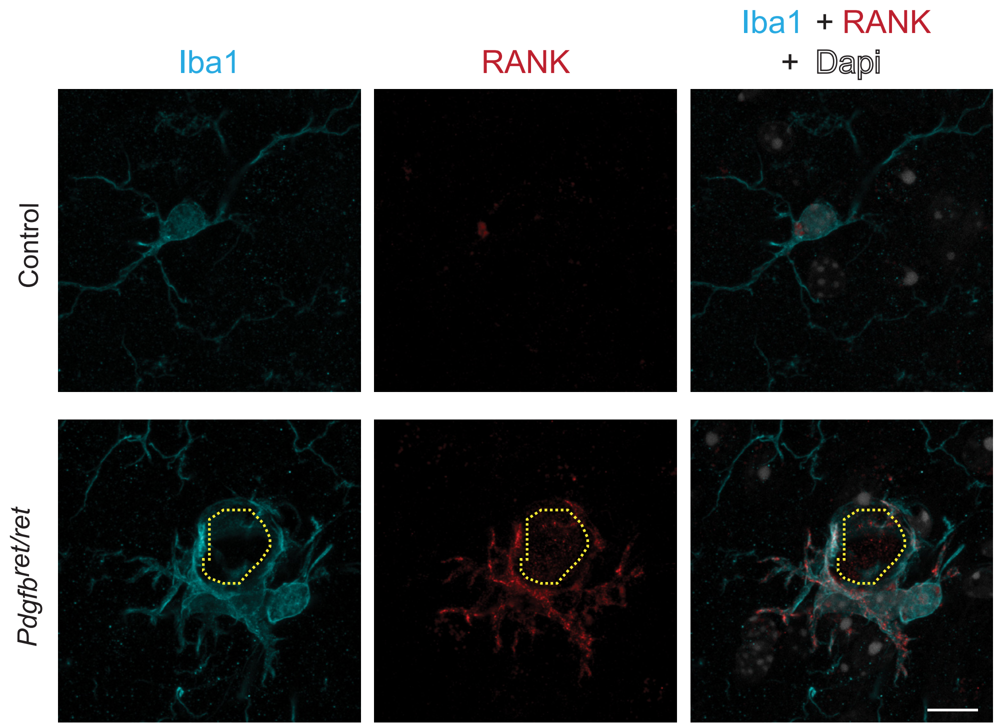

**Supplementary figure 5. Calcification-associated microglia express RANK.**

Iba1-positive (in cyan) microglia around brain calcifications (encircled with yellow dotted line) express the osteoclast marker RANK (red). Nuclei were visualized using DAPI (in white). Scale bar – 10 µm.

### Supplementary Figure 6.

Zarb et al.

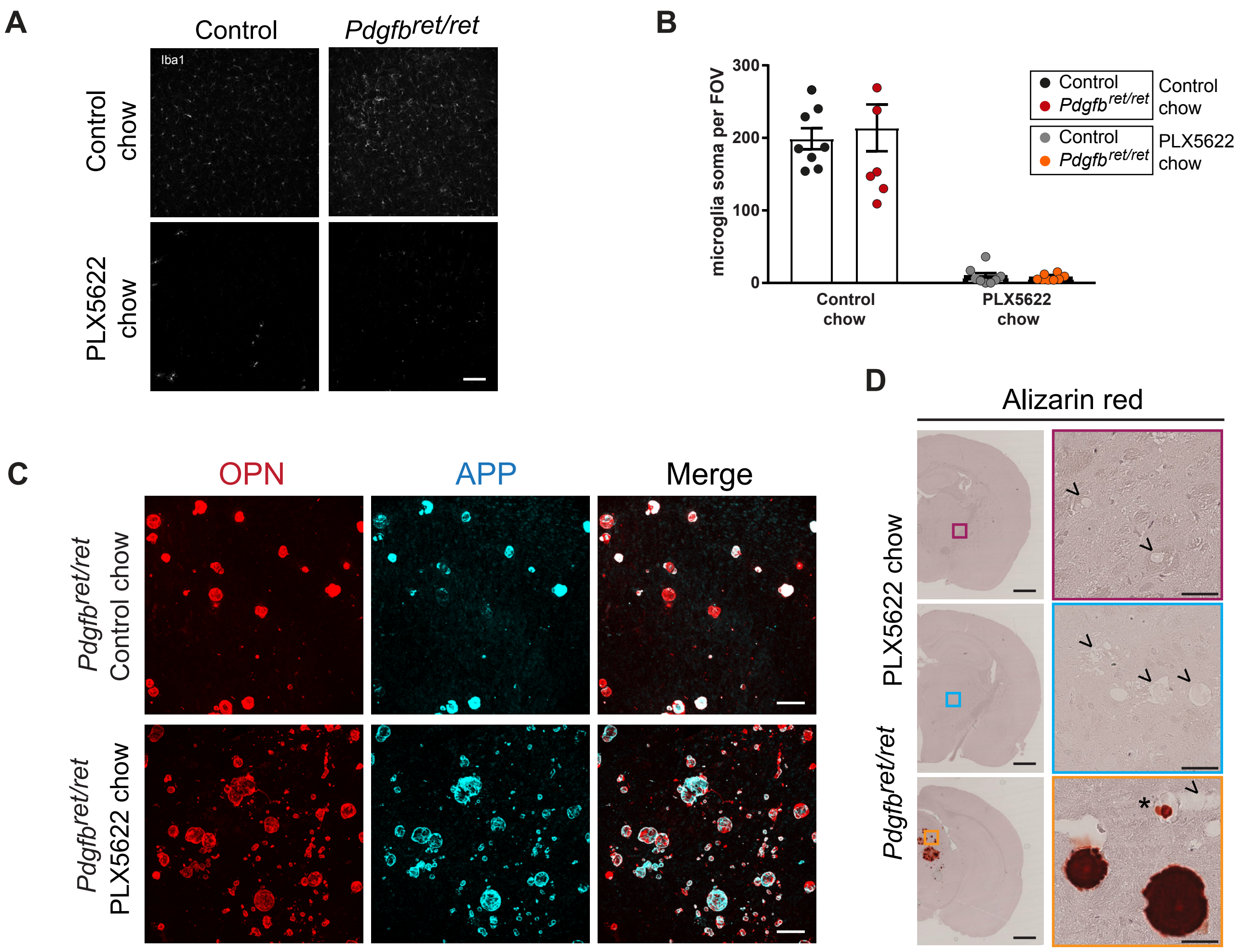

**Supplementary figure 6. Quantification of microglia, detection of OPN deposition and alizarin red staining of brain calcifications after PLX5622 treatment. (A)** Microglia (Iba1, in white) numbers are reduced both in *Pdgfb*$^{ret/ret}$ and control mice at 3 months of age after administering PLX5622 for two months. **(B)** Quantification of microglia shows a reduced number in *Pdgfb*$^{ret/ret}$ and control after PLX5622 administration. **(C)** OPN (in red) is deposited within brain calcifications in *Pdgfb*$^{ret/ret}$ animals (APP, in cyan) after PLX5622 treatment (lower panel). **(D)** Alizarin red staining of brain sections of *Pdgfb*$^{ret/ret}$ mice treated with PLX5622. Inclusions (indicated by arrowheads) in white matter (upper panel, in pink) and in the thalamus (middle panel, in blue) are negative. Some inclusions in the midbrain are positive for Alizarin red (in red), some inclusions are partially positive (indicated by an asterix) and others negative (indicated by an arrowhead). Scale bars –100 μm (A, C) and 1000 μm (D) and 50 μm (D inset).

### Supplementary Figure 7.

Zarb et al.

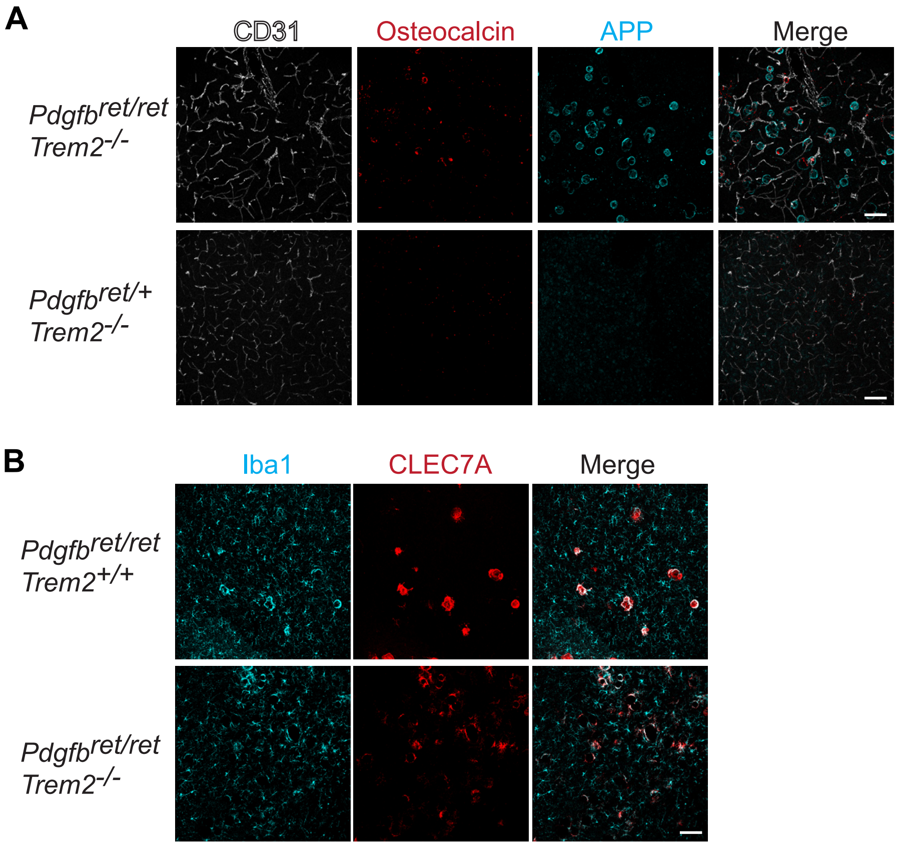

**Supplementary figure 7. Microglial activation and brain calcifications in a *Trem2*$^{-/-}$ background.**

**(A**) Brain calcifications are present in *Pdgfb*$^{ret/ret}$*; Trem2*$^{-/-}$ but not in *Pdgfb*$^{ret/+}$*; Trem2*$^{-/-}$ animals. Calcifications are visualized using osteocalcin (in red) and APP (in cyan), and blood vessels are visualized using CD31 (in white) immunohistochemistry. **(B)** Calcification-associated microglia (Iba1, in cyan) in *Pdgfb*$^{ret/ret}$ mice express the DAM-signature protein, CLEC7A (in red), in the presence and absence of *Trem2*. Scale bars – 100 µm.
