## Supplementray Movie 1 legend for "Microglia control small vessel calcification via TREM2"

**Supplementary video 1. Osteocalcin-positive striped structures in the white matter of mice administered with PLX5622 are not vessel-associated.** Vessels are visualized using CD31 (in white), with osteocalcin staining in red. A surface mask is applied for better visualization.
