## Supplementary Table 1 for "Microglia control small vessel calcification via TREM2"

**Supplementary table1. Primary antibodies used for immunohistochemistry.**

| **Antibody** | **Company** | **Cat. No.** | **Dilution** |
|---|---|---|---|
| Goat anti-osteocalcin | ABDserotec | 7060-1815 | 1:300 |
| Goat anti-osteocalcin | Genetex | GTX39510 | 1:500 |
| Goat anti-osteocalcin | Alfa Aesar | J65216 | 1:500 |
| Rabbit anti-Cathepsin K | Abcam | ab19027 | 1:100 |
| Rabbit anti-Collagen IV | ABDserotec | 2150-1470 | 1:300 |
| Rat anti-GFAP | Invitrogen | 13-0300 | 1:200 |
| Rabbit anti-GFAP | DAKO | Z0334 | 1:100 |
| Goat anti-podoplanin | R&D systems | AF3244-SP | 1:100 |
| Goat anti-TIMP2 | R&D systems | AF971 | 1:100 |
| Rat anti-CD68 | BioRad | MCA1957 | 1:600 |
| Rat anti-CD45 | BD Pharmingen | 553076 | 1:100 |
| Rabbit anti-APP | ThermoFischerScientific | PA5-19923 | 1:200 |
| Rat anti-CD31 | Dianova | DIA-310 | 1:100 |
| Rabbit anti-Ki67 | Thermo Fischer | MA514520 | 1:20 |
| Rat anti-PDGFRB | eBioscience | 14-1402 | 1:50 |
| Rabbit anti-Iba1 | WAKO | 019-19741 | 1:1000 |
| Goat anti-Iba1 | Abcam | ab5076 | 1:500 |
| Rabbit anti-ASC | Adipogen | AG-25B-0006 | 1:500 |
| Rat anti-RANK | Lifespan Bioscience | LS-C150237 | 1:100 |
| Goat anti-OPN (*Spp1*) | R&D systems | AF808 | 1:100 |
| Goat anti-LCN2 | R&D systems | AF1857 | 1:100 |
| Rat anti-CLEC7A (Dectin1) | Invivogen | Mabg-mdect | 1:30 |
| Rat anti-C3 | Abcam | ab11862 | 1:100 |
| Chicken anti-MBP | Millipore | AB9348 | 1:100 |
