## Supplementary Table 2 for "Microglia control small vessel calcification via TREM2"

**Supplementary table 2. Antibodies used for flow cytometry analysis.**

| **Antibody/reagent** | **Company** | **Cat. No** |
|---|---|---|
| anti-mouse CD206 (clone C068C2) conjugated with Alexa Fluor 700 | BioLegend | 141734 |
| anti-mouse CD11c (clone N418) conjugated with PE-Cy5.5 | eBioscience | 35-0114-82 |
| anti-mouse CD11b (clone M1/70) conjugated with Brilliant UltraViolet 737 | BD | 564443 |
| anti-mouse MerTK (clone DS5MMER) conjugated with PE-Cy7 | eBioscience | 25-5751-82 |
| anti-mouse Ly6C (clone HK1.4) conjugated with Brilliant Violet 711 | BioLegend | 128037 |
| anti-mouse Ly6G (clone 1A8) conjugated with Brilliant UltraViolet 563 | BD | 565707 |
| anti-mouse Lyve1(clone ALY7) conjugated with eFluor 660 | eBioscience | 50-0443-82 |
| anti-mouse CD38 (clone 90) conjugated with Alexa Fluor 488 | BioLegend | 102714 |
| anti-mouse I-A/I-E (clone M5/114.15.2) conjugated with Brilliant Blue 700 | BD | 746197 |
| anti-mouse Ki67 (clone 16A8) conjugated with BV421 | BioLegend | 652411 |
| anti-mouse F4/80 (clone BM8) conjugated with Brilliant Violet 510 | BioLegend | 123135 |
| anti-mouse CD64 (clone X54-5/7.1) conjugated with BV605 | BioLegend | 139323 |
| anti-mouse CD45 (clone 30-F11) conjugated with Brilliant UltraViolet 395 | BD | 565967 |
| anti-mouse CD14 (clone rmC5-3) conjugated with PE | BD | 553740 |
| anti-mouse CCR5 (clone C34-3448) conjugated with Biotin | BD | 559922 |
| Streptavidin PE-Cy5 | Biolegend | 405205 |
